# A CRISPR-Cas9 Genome Engineering Platform in Primary CD4+ T Cells for the Interrogation of HIV Host Factors

**DOI:** 10.1101/205500

**Authors:** Judd F. Hultquist, Joseph Hiatt, Kathrin Schumann, Michael J. McGregor, Theodore L. Roth, Paige Haas, Jennifer Doudna, Alexander Marson, Nevan J. Krogan

## Abstract

CRISPR-Cas9 gene editing strategies have revolutionized our ability to engineer the human genome for robust functional interrogation of complex biological processes. We have recently adapted this technology to primary human T cells to generate a high-throughput platform for analyzing the role of host factors in pathogen infection and lifecycle. Here, we describe applications of this system to investigate HIV pathogenesis in CD4+ T cells. Briefly, CRISPR-Cas9 ribonucleoproteins (crRNPs) are synthesized *in vitro* and delivered to activated primary human CD4+ T cells by nucleofection. These edited cells are then validated and expanded for use in downstream cellular, genetic, or protein-based assays. Our platform supports the arrayed generation of several gene manipulations in only a few hours’ time and is widely adaptable across culture conditions, infection protocols, and downstream applications. We present detailed protocols for crRNP synthesis, primary T cell culture, 96-well nucleofection, molecular validation, and HIV infection with additional considerations for guide and screen design as well as crRNP multiplexing.

## INTRODUCTION

Viral pathogens depend on the specialized microenvironments of their hosts for optimal replication and transmission. As obligate intracellular parasites, viruses rely on host proteins or ‘dependency factors’ to successfully replicate and infect new cells. In turn, the host has evolved molecular defenses known as ‘restriction factors’ to interfere with the replicative cycle of pathogens^1–3^. Agents that specifically target the interface of viral and host proteins are promising candidates for the development of next-generation therapeutics. As such, much effort has been expended on identifying and characterizing the molecular basis of these interactions. However, the tension between the fidelity of a model – the degree to which the experimental system recapitulates and identifies authentic *in vivo* relationships – and its genetic tractability and scalability has thus far compromised our ability to investigate these crucial interactions in a systematic and representative way.

The study of human host-pathogen interactions relies on the development and experimental manipulation of animal models *in vivo*, cell line models *in vitro*, and primary cell samples *ex vivo*. While appropriate animal models remain the gold standard for studying systemic pathogenesis, limitations in the genetic tractability, affordability, scalability, and ease of manipulation often demand the use of alternate model systems^4–6^. Conversely, while most *in vitro* cell line models are readily scalable and tractable, they often fail to recapitulate the relevant cellular and organismal microenvironments that exist during natural infection^7–9^. The use of immortalized human cell lines, for example, has yielded an extraordinary wealth of knowledge concerning host-pathogen relationships, but many of the findings extracted from these systems fail to recapitulate *in vivo*^10–12^. The process of immortalization, selection, and expansion of these lines often dramatically changes cellular expression profiles and responses to complex stimuli like infection^13,14^. This is especially true of genes involved in cellular metabolism, cell division, and signal transduction^14,15^. These experimental concerns and the limited translational capacity of the resultant findings have driven interest in primary cell models of disease, but primary cells are often difficult to obtain, culture, and manipulate *ex vivo*. Only recently have advances in genome engineering made primary cells a tractable genetic system *ex vivo*, allowing for a model system that combines the tractability of *in vitro* systems with the fidelity of *in vivo* models^16,17^.

Unbiased genetic screening approaches to uncover host-pathogen interactions – previously possible only in cell-line-based model systems – have been further complicated by limitations in tools for functional genetic perturbations. Previous genetic screening methods to define host-pathogen relationships relied heavily on RNA interference (RNAi) to alter gene expression^8,18–20^. While RNAi gene knock-down methodologies have provided an invaluable tool to biologists, they often suffer from low penetrance, transient efficacy, and high rates of off-target effects^7,21^. These concerns are particularly pronounced for gene products that either have a long half-life or are required at only low abundance (such as in the case of the retroviral integrase interactor, LEDGF ^22–25^). Even if a set of efficacious RNAi reagents are identified, validated, and stably integrated, knock-down of the gene product may be insufficient to reveal functional significance leading to low sensitivity^7,21^. Furthermore, off-target effects compromise the specificity of the screen and necessitate extensive validation of all hits^26,27^.

Due to these and other limitations to genetic manipulation and pathogen propagation *in vitro*, most attempts to comprehensively and systematically define host-pathogen interactions have yielded only a limited number of verifiable associations. Meta-analysis of three genome-wide RNAi screens for human-HIV host-pathogen interacting factors found a less than 7% overlap in candidates between any two studies and an overlap of only three genes between them all^7^. Likewise, meta-analysis of eight genome-wide RNAi screens for human influenza A virus (IAV) host-pathogen interacting factors found only a 7% overlap in candidates between any two studies^28^. In both cases, the variation between studies has been ascribed to differences in the RNAi libraries used to screen host candidate genes, the *in vitro* systems used to model infection, the readout of pathogen infectivity, and the strain of the pathogen itself^7,21,28^. While pathway and complex level analysis of these same datasets have revealed some novel insight^7,28^, new genetic tools and more functionally relevant models are required for the systematic identification and improved understanding of human host-pathogen relationships. Indeed, the development of new screening technologies and model systems in recent years has unveiled an array of new biology that continues to refine our understanding of viral infection and pathogenesis^9,29–32^.

The recent advent of CRISPR-Cas9 genome editing offers an alternate strategy for gene manipulation that improves upon key failings of previous approaches. Cas9 is a programmable DNA endonuclease that can be directed to a complementary region of the genome by its associated CRISPR (or guide) RNA^33–35^. Unlike RNAi approaches, which suppress messenger RNA, Cas9 targets the DNA directly for cleavage. Imperfect repair of the resultant double strand break can result in insertions or deletions (Indels) and lead to nonsense or frameshift mutations in coding regions, permanently ablating protein expression^36,37^. Alternately, concurrent delivery of a specific repair template can result in the precise excision or introduction of new sequences by homology-directed repair^17,33,34^. As these edits occur at the level of the DNA, they are heritable, stable, and completely penetrant, allowing for long-term expansion of populations without selection and complete clearance of even long-lived proteins^9,16^. Furthermore, the CRISPR-Cas9 editing has been shown to occur with higher efficiency and with fewer off-target effects than RNAi manipulation^38–42^. This highly precise, highly efficient, and permanent editing is in direct contrast with previous RNAi approaches. Employing this strategy in primary cells promises unprecedented fidelity and translatability of experimental results to native infection.

Here, we describe a protocol for high efficiency genome engineering in primary CD4+ T cells by CRISPR-Cas9 for the identification and exploration of human-HIV host-pathogen interactions (**Figure 1**). CRISPR-Cas9 ribonucleoproteins (crRNPs) are synthesized *in vitro* by the incubation of target-specific guide RNA, trans-activating crRNA (tracrRNA), and Cas9 protein^16,17^. These preformed complexes are then delivered to activated primary T cells by nucleofection for editing^16,17^. We have shown that this approach can be used to screen a wide array of genes for phenotypic influence with minimal manipulation, toxicity, or off-target impact. These reactions can be carried out in arrayed, 96-well format and can be readily multiplexed for the generation of double-knock-out pools. While we explore the use of this platform for the identification of human-HIV host-pathogen interactions, this general strategy can be applied more broadly to other primary cell types for the interrogation of a broad array of complex biological phenomena directly in human patient and donor samples.

**Figure 1.**
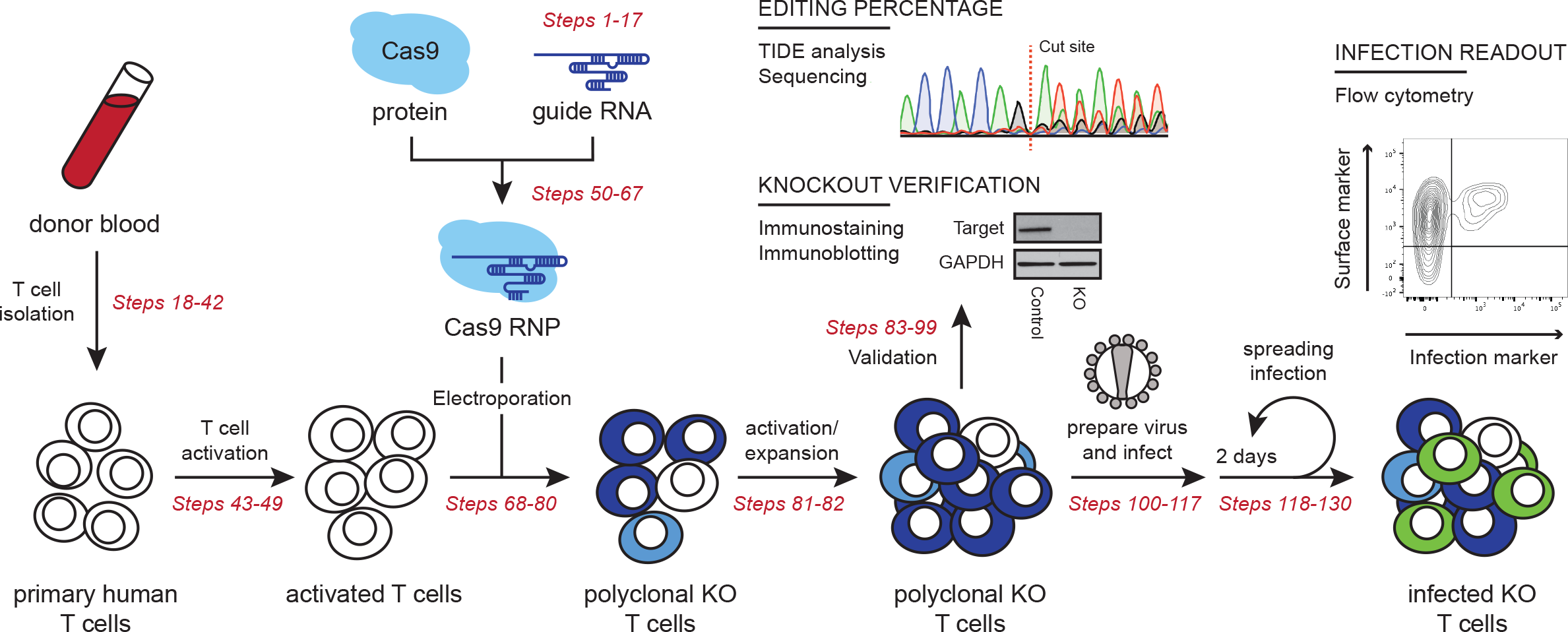
Experimental overview of primary T cell editing using CRISPR-Cas9 RNPs. Primary CD4+ T cells are isolated from donor blood and activated. crRNPs are synthesized *in vitro* and delivered to the activated T cells by nucleofection. These cells are expanded for molecular validation of gene editing and downstream phenotypic assays. HIV-1 virus stocks are prepared and used to infect the cell pools, which are monitored for infection by flow cytometry over several days. Genes that significantly alter infection relative to non-targeting controls are potential host factors for further mechanistic studies.

### Applications of crRNP-mediated Primary Cell Gene Editing

We have previously demonstrated that crRNPs can be successfully used to identify and screen new HIV host factors directly in primary CD4+ T cells^16^. We have furthermore demonstrated that this technology can be used to validate the findings of hypothesis and discovery-driven efforts in cell line models^9,16^. Given the scalability and ease of use, we believe that this approach is well suited for broad application not only to the study of host-pathogen interactions, but other T cell processes such as autoimmunity and immune regulation, pathogen sensing, and immunologic memory. Furthermore, crRNP multiplexing will allow for the study of epistatic relationships and functional redundancy among gene families. Delivery of these same crRNPs to other primary cell types has the potential to transform many such cells into tractable genetic models with numerous applications across basic, translational and clinical research.

The protocol described here has been successfully employed for screening potential host factors that interact with HIV Integrase to influence HIV infection^16^. We are currently expanding these efforts to screen thousands of genes in parallel. Simultaneously, we are working to enhance multiplexing efficiency to perform systematic epistasis mapping directly in primary cells. We are furthermore optimizing delivery to alternate primary cell types in an effort to study regulatory differences in different cellular subsets from the same donor. Helpful tips gleaned from these efforts have been included in the text where applicable to assist in the optimization of these approaches to additional fields of study.

### Limitations of crRNP-mediated Primary Cell Gene Editing

While broadly applicable, scalable, and tractable, this approach carries a number of limitations. First, delivery of *in vitro* synthesized crRNPs does not innately include a genetic-based selection marker. This lack of selection and the imperfect repair of the generated double strand breaks after cutting results in a polyclonal population of cells including unedited, heterozygous knock-out, and homozygous knock-out cells with heterogeneous sequences at the target locus. In other words, unlike RNAi, where a given reagent results in broad-based knock-down of protein expression across a population of cells, CRISPR-Cas9 editing will result in complete ablation of protein expression in only a certain proportion of cells. Phenotypic readout, therefore, is directly linked to the efficiency of the guide RNA and the resultant frequency of these populations in the final pool^16,17^. While non-ideal for phenotypic readouts that depend on the homogeneity of a cellular population, the all-or-nothing nature of these cells allows for the detection of phenotypes for proteins required at only a fraction of their cellular levels and with guide RNA of sub-par efficiency.

Fortunately, many loci are amenable to high efficiency editing, resulting in near complete ablation of protein expression in a cellular population even without selection. The guide RNA design protocols detailed here will typically yield at least one guide out of every three with an editing efficiency greater than 80 percent^16,39,43^. Some loci, however, appear refractory to CRISPR-Cas9 editing independent of guide design. While we hypothesize this may be due to the genome architecture or chromatin structure at the locus, it is unclear why this is the case^44,45^. Similarly, certain cell types appear generally more resistant to editing than activated T cells independently of the efficiency of crRNP delivery. Regardless of the cause, this underscores the importance of validating editing efficiency in every experiment and with every donor.

Second, while delivery of preformed crRNPs has yielded high efficiency editing in primary cell types, the transient lifespan of these complexes limits the potential of this delivery method for applications beyond gene editing. Notably, CRISPR interference (CRISPRi) and CRISPR activation (CRISPRa) approaches rely on continual occupancy of target loci by the Cas9 RNP complex^46–49^. Therefore, one-time delivery of preformed CRISPRi or CRISPRa RNPs would not be expected to cause long-term perturbations to gene expression, though transient perturbations may be achievable.

Third, as gene ablation is complete in successfully edited cells, the editing of essential genes leads to rapid cell arrest or death and the preferential expansion of unedited cells in the population. Thus, verification of editing efficiency in the population should be performed on cellular lysates harvested at the time of phenotypic analysis. Below we describe optional methods to verify editing at the protein level by immunoblotting or immunostaining and at the genetic level by DNA sequencing.

## EXPERIMENTAL DESIGN

### Considerations for Guide Design

Since the advent of CRISPR-Cas9 editing, several different protocols and algorithms for the design of guide RNA have been described (*i.e*., ^39,50,51^, reviewed in^52^). These algorithms have improved markedly over time, resulting in higher on-target hit rates with fewer off-target effects^52,53^. At the time of writing, the best such algorithm is arguably Rule Set 2^39^ A number of resources exist to identify the best guide RNA sequences for a given gene, a few of which are described below. While we design and order our custom guide RNAs for synthesis by contract, they can also be generated by *in vitro* transcription or can be ordered pre-designed from a number of commercial providers^16,17^. If using pre-designed guides, we recommend checking the specificity of each guide *in silico* prior to purchase to ensure it targets the proper locus with minimal affinity for other genomic sequences. Stability modifications to the RNA guides (such as 2xMS) can be added at no loss to editing efficiency (*e.g*., ^54^).

For the generation of gene knock-outs, we recommend ordering three to five distinct guide RNAs per gene. For most loci, this will yield at least one guide that exhibits high efficiency editing. While any part of the gene can be targeted for editing, it is essential to take the gene structure into account during the design process to avoid intron-exon boundaries and alternatively spliced sites. Many genome browsers will provide isoform information to assist in targeting key exons or, conversely, for targeting only those forms of the protein you wish to ablate. We describe Ensembl as one example below, but we have had success with a variety of online tools. Benchling, for example, is free for academic use, and provides an online tutorial for guide design at https://benchling.com/tutorials/21/designing-and-analyzing-gRNAs. Furthermore, recent publications have provided pre-designed guides for human and mouse genomes^39^, many of which are available through Addgene.

### Considerations for Primary Cell Isolation and Culture

Unlike cell lines, which can be expanded indefinitely, primary cells have limited longevity and expansion potential, so isolation of a sufficient number of cells is of paramount importance. The protocols detailed here have been verified to be effective for inputs of 200,000 to 1,000,000 cells per nucleofection reaction, and may be applicable for cell counts both below and above this range with some loss to editing efficiency. Cells will continue to expand after editing and can be maintained in culture for several weeks post-nucleofection. Any scalable T cell isolation protocol may be used to procure the required number of cells from whole blood or blood product and many robust, compatible protocols exist. We describe isolation of peripheral blood mononuclear cells (PBMCs) from whole blood and the subsequent isolation of CD4+ T cells by negative selection using the StemCell EasySep T Cell Isolation Kit.

Regarding CD4+ T cell activation, we have achieved successful editing after stimulation with either plate-bound anti-CD3 and soluble anti-CD28, bead-bound anti-CD3/anti-CD28, bead-bound anti-CD2/anti-CD3/anti-CD28, or PHA and IL-2. When considering stimulation, we find there is a balance between the viability of cells after editing, which is promoted by gentler stimulation, and editing efficiency, which is promoted by stronger stimulation. Therefore, for knock-out experiments like those described here, we recommend a gentle stimulation protocol with plate-bound anti-CD3 and soluble anti-CD28 for 72 hours prior to nucleofection, followed by a second round of stronger stimulation with bead-bound anti-CD2/anti-CD3/anti-CD28 immediately after nucleofection to maximize cell proliferation. The purity and activation state of the CD4+ T cells should be monitored by CD4 and CD25 staining on the day of nucleofection.

If it is anticipated that knock-out of targeted genes will alter cell growth phenotypes, we recommend proceeding with assays three to seven days post-nucleofection. This allows enough time for the cells to recover from the reaction and for the existing protein to degrade, but not so much time that unedited cells can outcompete or dilute out edited cells in the culture.

### Considerations for Screening

Screen design is closely tied to the number of genes being screened and the downstream phenotypic assays. For targeted validation of a few genes, we recommend validating guides with a test batch of primary T cells before embarking on phenotypic analysis. Guide efficiency varies by sequence, target locus, and cell type, but appears relatively consistent between donors. Indeed, while we observe donor-to-donor variation in the overall permissivity to CRISPR-Cas9-mediated gene editing, the relative efficiency of editing between different guide RNAs is nearly always consistent between donors.

For medium-to-large-scale screens with three to five guides per gene, it may be easier to embark on the phenotypic assays without prior guide validation. Guide RNAs can be ordered from commercial sources pre-arrayed in 96-well plates. Conveniently, this orientation may be preserved for crRNP generation, nucleofection into cells for editing, HIV spreading infections, and additional other phenotypic assays. Genomic or protein lysates can also be stored in this format for later validation. Thus, it may be expedient to proceed with phenotypic assays after validation of the positive controls only, limiting broad validation of guide RNA editing efficiencies to only those guides that display a significant phenotype. One consequence of this strategy is that “negative” results in phenotypic assays are difficult to interpret; they may indicate that the target host factor has no role in the process under examination or they may represent editing failures. Disambiguation requires validation of every guide, which may or may not prove worthwhile depending on the application and experimental design.

### Controls

As with any experimental system, the right controls are critical for downstream interpretation of results. The importance of such controls grows in proportion to the size of the experiment as variations in donors, plate handling, and environment must be robustly accounted for. While the controls we recommend below are appropriate for infection-centric studies of HIV host-pathogen interactions, additional controls appropriate to the specifics of the downstream phenotypic assay are strongly recommended. Generally, each plate of a screen or each small experiment should have at least three types of controls: a negative, non-targeting control; a positive control with a strong effect on the phenotype being assayed; and a toxicity control.

An important negative control for all applications is nucleofection of a crRNP that bears a non-targeting guide RNA. Several such sequences have been published and still more are commercially available as proprietary reagents^39,50,55^. We recommend including at least two distinct non-targeting guides per experiment, randomly distributed if on a plate. Other negative controls, including unedited cells, cells nucleofected with nothing, or cells nucleofected with Cas9 protein alone may or may not be appropriate for the assay at hand.

We also recommend at least two distinct positive controls that have predictable effects on the phenotype being assayed. For HIV replication, we have found that the dependency factors CXCR4 and LEDGF are both effective and reproducible controls (see guide RNA sequences below). These guides result in the near complete ablation of protein expression and inhibit HIV infection to different degrees by different mechanisms. *CXCR4* knock-out prevents HIV entry into cells in a manner dependent on virus tropism and consistently decreases infection rates to near the limit of detection^16^. *LEDGF* knock-out, on the other hand, partially inhibits proviral integration and inhibits infection by typically 50-75%, but in a tropism-independent manner^16^.

One final control we recommend is knock-out of a gene essential to cell health and division, such as *CDK9*. In most donors, successful knock-out results in significant toxicity and cell death. Comparison of this well’s cell count, viability, and phenotype to that of experimental wells is useful for setting thresholds for analysis and for identifying other potentially essential genes. While the toxicity of gene knock-out will vary dependent on cell type, culture conditions, and timing, several lists of essential human genes have been previously published and may be indicative of potential toxicity^56–58^.

### Considerations for Viral Replication

One or more viruses may be pertinent for downstream analyses. Generally, a molecular clone with a reporter gene is strongly recommended for ease of downstream processing. The use of a fluorescent reporter obviates the need for staining of cells before flow cytometry analysis, saving reagents, time and the multiple centrifugations that can quickly become overwhelming when a large number of plates are in culture. In the protocol below, we use the replication-competent HIV-1 clone NL4-3 Nef:IRES:GFP^59^. The virus can be prepared in bulk and concentrated for use across multiple experiments. Viruses with different reporters (such as luciferase^60^), may be used alternately depending on the preferred readout. While we titer each new preparation of virus against primary CD4+ T cells, note that each donor will vary in his or her susceptibility to infection and therefore live virus titers may vary significantly from one donor to the next. To prevent repeated live virus titering, one alternative is to calculate p24 concentration and infecting at equal mass across donors. If high rates of infection are required, spinoculation of cultures may be required to boost initial infection rates^61^.

### Consideration for studies in other cell types

The protocol reported here has been optimized for use with CD4+ T cells but can be adapted successfully to other T cell subsets as well as other primary cell types. The key hurdle to adapting this protocol for additional cell types is optimization of crRNP delivery, though additional variables may also influence successful editing including chromatin architecture, cell viability, and activation state. Successful delivery of the crRNPs may be monitored by immunostaining for Cas9 after nucleofection or by nucleofection of a fluorescently labeled Cas9 protein, but this may be difficult depending on the cells in question. We therefore typically recommend optimizing by quantifying editing efficiency at the genomic and/or protein levels under various conditions using previously validated guides. As guide efficiency is not guaranteed to be consistent between cell types, we recommend doing these optimization experiments with at least two independent guides.

When optimizing delivery to a new cell subset, pay special attention to input cell stimulation and concentration, cell culture conditions, nucleofection buffer, and nucleofection pulse code. The confluence of these factors will impact both editing efficiency and cell viability, collectively determining the frequency of edited cells in your resultant population. Several nucleofection optimization kits designed to test a range of conditions and buffers are commercially available (see, for example, Lonza catalog number V4XP-9096) and new delivery strategies for various cell types are published regularly (*e.g*., ^17,36,43,62^). The development of these strategies for peripheral and tissue-localized hematopoietic cell subsets, epithelial tissues, and an array of progenitor cells including induced pluripotent stem cells (iPCs) promises to revolutionize genetic study of primary human samples for the advancement of a wide array of disease research.

## MATERIALS

### Reagents

Whole human blood or leukoreduction chamber
Ficoll-Paque (Fisher, cat. no. 17144003)
HEK293T cells (ATCC, cat. no. CRL-3216) for packaging of lentivirus
Fetal Bovine Serum (FBS) (Life Technologies, cat. no. 26140-079)
DMEM, high glucose (Corning, cat. no. MT10-017-CV)
RPMI-1640, high glucose (Corning, cat. no. MT10-040-CV)
100× penicillin-streptomycin (Corning, cat. no. MT30-002-CI)
100× sodium pyruvate (Corning, cat. no. MT25-000-CI)
100× Hyclone HEPES (Fisher, cat. no. SH3023701)
PBS (Corning, cat. no. MT46-013-CM)
EDTA (Corning, cat. no. MT46-034-CI)
Bleach
70% ethanol (VWR, cat. no. 89125-172)
Milli-Q purified or cell-culture grade H2O
RNaseAway (Sigma, cat. no. 83931-250ML)
IL-2 (Miltenyi Biotec, cat. no. 130-097-744)
T Cell Activation and Stimulation Kit (Miltenyi, cat. no. 130-091-441)
T Cell Isolation Kit (StemCell Technologies, cat. no. 19052)
anti-CD3 clone UCHT1 (Tonbo, cat. no. 40-0038-U500)
anti-CD28 clone 28.2 (Tonbo, cat. no. 40-0289-U500)
CD4-PE, human (Miltenyi Biotec, cat. no. 130-091-231)
CD25-APC, human (Miltenyi Biotec, cat. no. 130-101-435)
Trypsin (Corning, cat. no. MT25-052-CI)
MACS Buffer (2% FBS, 2mM EDTA, PBS)
PolyJet (SignaGen Laboratories, cat. no. SL100688)
HIV-1 NL4-3 Nef:IRES:GFP full molecular clone (ARRRP, cat. no. 11349)
50% sterile polyethylene glycol 6000 (Fisher, cat. no. 528877-1kg)
4M NaCl, Sterile (Fisher, cat. no. BP358-212)
Synthetic crRNA (Dharmacon, cat. no. crRNA-XXXXX)
Synthetic tracrRNA (Dharmacon, cat. no. U-002000-50)
Purified Cas9-NLS protein (MacroLab, cat. no. Cas9-NLS)
Primary Cell Nucleofection Kit P3, Amaxa (Lonza, cat. no. V4SP-3096)
10mM Tris-HCl pH 7.4, RNAse-free (Corning, cat. no. MT46-030-CM)
2M KCl (ThermoFisher, AM9640G)
10mM dNTPs (NEB, cat. no. N0447L)
Phusion high-fidelity (HF) PCR kit with 5× HF buffer (NEB, cat. no. E0553L)
10mM Oligonucleotides for TIDE analysis (IDT, target specific sequences)
Formaldehyde (Sigma, cat. no. F8775-4X25ML)
2.5x Laemmli Sample Buffer (Sigma, cat. no. S3401-10VL or equivalent)
QuickExtract (EpiCentre, cat. no. QE09050)
DMSO (Sigma, cat. no. D2650-100ML)

### Equipment

Amaxa 4D Nucleofector with X Unit (and 96-well shuttle) (Amaxa, cat. no. V4XC-1032)
Tissue culture plates, 15 cm (Fisher, cat. no. 12-565-100)
Mr. Frosty Freezing container (Fisher, cat. no. 15-350-50)
T150 flasks (Fisher, cat. no. 1012634)
Single and Multichannel Pipetman
Hemacytometer (or equivalent cell counter)
Pipet-Aid
Flow Cytometer (ThermoFisher Attune NxT or equivalent)
48-well plates (Fisher, 08-772-52)
Non-treated 48-well tissue culture plates
1.5mL Cryotubes
PCR plates, 96-well (Bio-Rad, cat. no. MLP-9601)
Low-Bind 96-well plate (E&K Scientific, cat no 651261)
Aluminum plate seals (Bio-Rad, cat. no. MSF1001)
Falcon conical tubes, 15 and 50 ml (Fisher, cat. no. 14-959-53A and 14-432-22)
Invitrogen Xcell SureLock mini-cell (Invitrogen, cat. no. EI0001) or similar
Tissue culture benchtop centrifuge with plate buckets and biosafety caps
0.22μm Steriflip filters (Fisher, cat. no. SCGP00525)
Thermal cycler (Bio-Rad, cat. no. 170-9713)
Computer

### Reagents Setup

#### DMEM media

To 450mL of high glucose DMEM, add 50 ml of FBS (10% vol/vol) and 5 ml of penicillin-streptomycin (100×). Store the medium at 4°C for up to 3 months.

#### Freezing medium

Prepare freezing medium by adding 5 mL of DMSO to heat-inactivated FBS (10% vol/vol); filter it through a 0.2μm steriflip filter. Store the medium at 4 °C for up to 1 year.

#### Complete RPMI media for primary CD4+ T cells (cRPMI)

To 450mL of high glucose RPMI-1640, add 50mL of Fetal Bovine Serum (FBS, 10% vol/vol final), 5mL of penicillin-streptomycin (Pen/Strep, 50ug/mL final), 5mL of sodium pyruvate (NaPy, 5mM final), and 12.5mL of 4-(2-hydroxyethyl)-1-piperazineethanesulfonic acid (HEPES, 5mM final). Store the medium at 4°C for up to 6 months.

#### IL-2

Resuspend lyophilized IL-2 in Milli-Q purified H2O at 20IU/uL. Make 50uL aliquots in 500uL lo-bind tubes and store at −20 °C for up to 1 year.

#### MACS Buffer

Add 10mL FBS and 2mL 0.5M EDTA pH8.0 into a final volume of 500mL 1xPBS. Filter sterilize.

#### 2.5x Laemmli Sample Buffer

Dilute 1.87mL 0.5M Tris-HCl pH 6.8, 6mL 50% glycerol, 3mL 10% SDS, 750uL 2-mercaptoethanol, and 150uL 1% bromophenol blue to 30mL total volume with 1xPBS.

#### Guide RNA/Cas9 Resuspension Buffer

To 45.5mL of water add 500μL of 1M Tris-HCl for a final concentration of 10mM, add KCl to a concentration of 150mM, adjust pH to 7.5, then filter sterilize.

#### Stimulatory Beads

Prepare and store in accordance with the manufacturer’s instructions.

### Equipment Setup

#### Nucleofection Parameters

On the Amaxa 4D-Nucleofector, prepare a custom protocol for pulse code EH-115 with buffer P3 according to the manufacturers instructions.

## PROCEDURE

**Designing CRISPR Guides**, Timing 1d

1. Open up the Ensembl human genome browser (http://ensembl.org) and search for the gene symbol or Ensembl ID of your gene of interest.
2. Open the ‘Human Gene’ page associated with your gene. Near the top of the page it will list known transcript variants. TSL:1 variants have the best support. For a knock-out of all known isoforms, identify an exon shared by at least all TSL:1 variants.
3. Open the page for an isoform Transcript ID by clicking on the hyperlink.
4. ‘About this transcript’ will list the number of exons as a hyperlink. Click the link to display the exon sequences below.
5. For the best results, conserved exonic sequences immediately following the start codon should be targeted. If there is limited exonic coding sequence before the next intron, skip to the next exon of at least 100 nucleotides.
6. Copy 100-250 nucleotides of sequence.
7. Open the CRISPR design webtool hosted by MIT (http://crispr.mit.edu/). Paste the sequence in the ‘sequence’ box and select ‘Unique Genomic Region’.
8. Enter your e-mail address and the gene name into the appropriate fields. Submit Query.
9. Guides will update as searched, listing specificity scores and off-target hits.
10. Copy the sequence for 2-4 of the best scoring guides to an excel file. Don’t include the PAM sequence! We recommend also recording the name of the exon searched, the target gene, the guide score, and any pertinent notes to the search (i.e. exon choice, etc.). **Critical Step**: We do not recommend ordering guides that score under 65. If ordering many guides, it may be worth searching several exons for the best possible guide RNA sequences.
11. Place synthetic guide RNA orders through Integrated DNA technologies, Dharmacon, or an alternate appropriate vendor. Some companies offer RNA stabilizing modifications (*i.e*. 2xMS from Dharmacon), all of which are compatible with this protocol. Alternately, procedures for *in vitro* transcription can be found here^17^. Note that while multiple vendors for tracrRNA and crRNA exist, tracrRNA is typically not compatible between vendors. **Resuspending CRISPR RNA**, Timing 1d **Critical Step**: tracrRNA and crRNA should not be suspended until immediately before use or should be stored in small aliquots in PCR strip tubes and thawed immediately prior to use. Thawed aliquots should not be frozen again. **Pause Point**: Store suspended stocks and working aliquots at −80°C. RNAs are stable at −80°C in this buffer for at least 6 months. **Critical Step**: Guide RNA and tracrRNA should be subjected no more than one freeze-thaw cycle. To avoid repeated freeze-thaw cycles, some consideration should be given to aliquoting strategy. If working at large scale, crRNPs should be generated in plate format upon initial resuspension of crRNAs (see “CRISPR-Cas9 RNP Generation Protocol: Medium-Large Scale in Plates” below). If resuspending a smaller number of guides in tube format, we recommend aliquoting to PCR strip tubes in 5uL aliquots.
12. Prepare an RNAse-free biosafety cabinet by wiping down the work surface, vortex, pipets, and racks with 70% ethanol followed by RNAse-Away. Prepare a tray of ice for working with suspended RNA. A tray of dry ice for lyophilized guides and frozen aliquots may be desirable depending on the size of the experiment.
13. Spin down the tubes or 96-well plate of lyophilized RNA to ensure the material is on the bottom of the tubes or wells.
14. Add sufficient RNAse-free 10mM Tris-HCl, 150mM KCl pH 7.4 to each tube or well for a final concentration of 160uM. For example, suspend 20nmol guide RNA in 125uL.
15. Seal the tubes or plate and vortex briefly to aid suspension.
16. Pulse spin the tubes or plate again to return RNA to bottom. Keep all suspended RNA on ice.
17. Using a p20 multichannel and sterile filter tips, make at least 5 aliquots of 5uL each in 0.5mL lo-bind tubes or PCR strip tubes. If working with 96-well plates, replica plate the guide RNA into at least 5 96-well plates, again 5uL per well. Seal each tube or plate securely and label with the guide/plate ID, description, date suspended, and concentration. Store aliquots at −80°C or proceed immediately with crRNP synthesis (Step 50). **PBMC Isolation**, Timing 1d **Caution**: All human blood samples must be handled according to approved biosafety protocols and regulations set by each individual lab and institution. Improper handling of samples may lead to exposure to blood borne pathogens. **Caution**: All human blood samples and data obtained from them must be handled according to approved biosecurity protocols for the protection of patient confidentiality and health information. **Timing**: Isolations are best performed on Monday or Tuesday to allow 72 hours for activation prior to nucleofection. Isolations should be done within 12 hours of drawing. **Timing**: A standard PBMC and T cell isolation from one donor will typically take between 4 and 5 hours. Make sure sufficient time is secured in the appropriate biosafety hood on the day of isolation.
18. Prior to isolation, coat wells of a sterile, non-treated 48-well tissue culture plate with anti-CD3 antibody for T cell stimulation.
  a. Each well will hold 1.25 million cells during stimulation and each nucleofection can tolerate as few as 200,000 to as many as 1 million cells. Plan to coat enough wells to stimulate sufficient cells for your experimental needs.
  b. To each well, add 150uL of 10ug/mL anti-CD3 (clone UCHT1) diluted in PBS. Incubate the plate for 2 hours at 37°C. Plate coating may also proceed for 8 hours at room temperature (RT) or overnight at 4°C. For these protocols, wrap the plate in parafilm to prevent evaporation.
  c. Note that other culture plates may be used if stimulation is scaled appropriately. The suggested stimulation and culture conditions described herein have been optimized for the longevity and viability of the culture, but plate-bound stimulation may be eschewed entirely in favor of bead-bound stimulation provided the beads are magnetically removed prior to nucleofection.
19. Once the donor samples have arrived, post any required biosafety notices around your workspace, record any donor information in your logs, and equip any required PPE.
20. Per donor, prepare 8 15mL conical tubes with 4mL Ficoll-Paque.
21. If working with donor blood, carefully cut the drainage and ventilation cords and drain the sample into a sterile 50mL conical tube. Dilute 1:1 with PBS and mix gently by pipet.
22. If working with a leukoreduction chamber, carefully cut the top cord leading into the conical chamber with scissors. Cut the bottom cord from the “peak” and set the conical chamber on top of a 50mL conical tube to drain by gravity flow. It should release 5-10mL of concentrated leukocytes. Dilute the blood with PBS-EDTA to a final volume of 65mL split between two 50mL conicals.
23. Dispensing slowly with a 10mL pipet, carefully layer 8 mL blood/PBS mixture onto the Ficoll-Paque in the 8 15mL conical tubes. A clear boundary should be evident.
24. Centrifuge to separate the blood components at 400xg for 30 minutes at RT. Make sure that the brake is turned off or to the minimum setting for this spin.
25. The blood will separate into the following layers from top to bottom: plasma, PBMCs (buffy coat), Ficoll, Granulocytes, Erythrocytes. Carefully aspirate off the plasma top layer to only a half centimeter above the buffy coat.
26. Using a 5mL pipet, slowly remove the buffy coat and transfer to a 50mL conical. To recover all the PBMCs, usually you will aspirate between 1-2mL including some remaining plasma and Ficoll below. Pool cells from the same donor in a single tube.
27. Fill to 50mL with PBS-EDTA and centrifuge at 300xg for 10 minutes at RT (the brake can now be reapplied).
28. Remove the supernatant. Note that it will be cloudy due to suspended platelets.
29. Resuspend cells in 50mL PBS-EDTA and centrifuge at 200xg for 10 minutes at RT. Remove the supernatant and suspend in 50mL PBS-EDTA.
30. Count the PBMCs using a hemocytometer or cell counter. You may have to dilute a sample of the PBMCs 1:10 to get an accurate count.
31. After counting, centrifuge at 300xg for 10 minutes at RT and remove supernatant.
32. Resuspend cells at 5*10^7 cells/mL in MACS buffer (1xPBS, 2mM EDTA, 0.5% BSA).
33. From each donor, remove 50uL to a 1.5mL centrifuge tube. Pellet the cells in a microcentrifuge by spinning at 400xg for 3 minutes. Remove the supernatant and resuspend in 200uL complete RPMI [cRPMI: high glucose RPMI-1640, 10% Fetal Bovine Serum (FBS), 50ug/mL penicillin/streptomycin (Pen/Strep), 5mM HEPES, 5mM sodium pyruvate (NaPy)] media in a 96-well round-bottom plate. These PBMCs will be stained as enrichment controls (Step 52, **Figure 3A**). **CD4+ T Cell Enrichment (EasySep)**, Timing 1d
34. T cell enrichment will occur in 14mL polystyrene tubes. Depending on your blood source, you may end up with many more cells than necessary for your experimental purposes; tailor your PBMC isolations for subsequent purifications according to yield. Extra PBMCs can be cryopreserved for other uses, disposed of in bleach or used for monocyte purification, infections, etc.
35. Aliquot up to 5mL of the PBMC resuspension into a sterile 14mL polystyrene tube.
36. Add 50uL/mL of negative selection antibody cocktail from the EasySep T Cell Isolation Kit to each tube. Mix gently, but completely, and incubate at RT for 5 minutes.
37. During the incubation, resuspend the RapidSphere magnetic beads from the EasySep T Cell Isolation Kit by vortexing vigorously for 30 seconds.
38. Add 50uL/mL RapidSpheres to each tube. Mix gently by pipet and incubate at RT for 5 minutes.
39. Place the sample (without the lid) onto the EasySep magnet and incubate at RT for 5 minutes.
40. Pick up the magnet and pour off the remaining cell suspension into a clean, labeled tube.
41. Combine cells from the same donor, mix, and count the enriched T cells using a hemocytometer or cell counter.
42. From each donor, remove 100uL to a 1.5mL centrifuge tube. Pellet the cells in a microcentrifuge by spinning at 400xg for 3 minutes. Remove the supernatant and resuspend in 200uL cRPMI in a 96-well round-bottom plate. This ‘CD4 Enriched’ fraction will be stained or CD4 and CD25 (Step 52, **Figure 3A**). **T Cell Activation and Expansion**, Timing 2-3d
43. Pellet the T cells at 300xg for 5 minutes. Remove the supernatant and resuspend in cRPMI plus 20U/mL IL-2 and 5ug/mL anti-CD28 (clone CD28.2) for a final concentration of 2.5x10^6 cells/mL.
44. Aspirate the antibody mixture from the previously prepared activation plate. Anti-CD3 should now be bound to the plate.
45. Add 500uL of suspended cells per well of a 48-well plate or 3-4mL of suspended cells per well of a 6-well plate and incubate for 48-72 hours at 37°C/5% CO2/humid/dark.
46. Resuspend and count the cells in a representative well from each donor. The cells should have begun to multiply and should have gone from being small and round to the larger, irregularly shaped morphology characteristic of activated T cells.
47. Suspend, pool, and count the cells. Pellet the cells at 400xg for 3 minutes and resuspend in cRPMI plus 20U/mL IL-2 at a final concentration of 2.5x10^6 cells/mL. Note that no antibody as added to the media here.
48. From each donor, remove 100uL to a 1.5mL centrifuge tube. Pellet the cells in a microcentrifuge by spinning at 400xg for 3 minutes. Remove the supernatant and resuspend in 200uL cRPMI in a 96-well round-bottom plate. This ‘Post-Stimulation’ fraction will be stained for CD4 and CD25 (Step 52, **Figure 3A**).
49. **Critical Step**: Stain the reserved PBMCs, CD4 Enriched T cells, and Post-Stimulation T cells with CD4-PE and CD25-APC according to the manufacturers instructions. The stimulated cells should be >90% CD4+CD25+ positive (See Anticipated Results, **Figure 3A**). If so, proceed with the nucleofection and/or infection. **? Troubleshooting** **CRISPR-Cas9 RNP Generation Protocol: Small Scale in Tubes**, Timing 1d
50. Work in an RNAse-free biosafety cabinet as described above. All tubes and pipette tips should be sterile and RNAse-free.
51. Set a heat block, incubator, or thermocycler to warm to 37°C.
52. Thaw an aliquot of guide and tracrRNA (160uM) and warm to RT.
53. Per nucleofection: Mix 1uL tracrRNA with 1uL crRNA. They will anneal to form 2uL of guide RNA at an effective concentration of 80uM. Be sure to generate crRNA for appropriate positive and negative controls at the same time (*i.e*. non-targeting guide, CXCR4, etc.; see **Controls** above).
54. Cap tubes and incubate RNAs together at 37°C for 30 minutes. (If using an incubator, it is advisable to increase this incubation time to 40 minutes to account for the slower temperature change of convection versus conduction.) Near the end of the incubation, thaw an aliquot of Cas9 protein (40uM) and warm to RT.
55. Add 2uL Cas9 to the RNA mixture by moving the pipette tip in small, rapid circles while ejecting VERY SLOWLY. The 2:1 molar ratio of guide RNA to Cas9 promotes efficient crRNP formation, yielding a preparation of crRNPs at 20uM. **Troubleshooting**: Rapid addition of the Cas9 may cause precipitation. If a precipitant forms, flick the tube or gently pipet up and down until the protein returns to suspension.
56. Incubate at 37°C for 15 minutes to form crRNPs. **Pause Point**: The steps outlined here generate sufficient crRNPs for a single reaction. crRNPs may be stored at −80°C in this buffer for at least 6 months or 4°C for up to two weeks with little-to-no loss in editing efficiency. Freeze/thaw of crRNPs is not advisable but limited cycles do not significantly impact editing efficiency.

#### BOX 1

##### Considerations for Scale and crRNP Multiplexing

Each nucleofection reaction is limited by the stable concentration of crRNPs in solution and the volume of crRNPs that can be added to the nucleofection cuvette without detrimental dilution of the nucleofection buffer. The volume of crRNPs can be increased up to twofold from what is recommended here without diminishing nucleofection efficiency. While this may provide increased editing efficiency for otherwise inefficient guides, most targets do not require this modification and the volumes recommended provide a good balance of editing efficiency, convenience, and cost at large scale. Given that the volumes can be effectively doubled in the reaction, two independent crRNPs can be delivered to the same set of cells simultaneously, either against the same or different genes. This allows for the generation of condensed screening platforms and even double knock-out populations for the study of epistatic relationships and functional redundancy^16^. For multiplexing, we recommend preparation of each pool of crRNPs independently with mixing of the pools at a 1:1 ratio prior to nucleofection. Delivery of more than two independent crRNPs per reaction is likely to come at an efficiency cost and may require additional optimization. **CRISPR-Cas9 RNP Generation Protocol: Medium-Large Scale in Plates**, Timing 1d
57. **Timing**: If working with crRNAs lyophilized in plate format, it is recommended to generate crRNPs at the time of guide RNA resuspension.
58. Work in an RNAse-free biosafety cabinet as described above. All tubes and pipette tips should be sterile and RNAse-free.
59. Set a 96-well heat block, incubator, or thermocycler to warm to 37°C.
60. Thaw plates of lyophilized guide RNAs and resuspend as above in 10mM Tris-HCl, 150mM KCl pH 7.4 to a final concentration of 160uM (Steps 12-20). Pulse spin the plate to ensure the RNA is at the bottom of the well.
61. Thaw aliquots of tracrRNA and pulse spin the tubes to ensure tracrRNA is at the bottom of the tubes.
62. In a separate PCR plate, aliquot 5uL 160uM tracrRNA to each well that will be used to generate guide RNA. Keep the PCR plate on ice. Note the volume added will depend on the amount of crRNP prepared. The instructions below generate enough crRNPs for five reactions.
63. Using a multichannel pipettor, add 5uL (or an equal volume, as appropriate) of resuspended crRNA from the source plate to the plate of tracrRNA. Seal the plate and incubate for 30min at 37°C to form guide RNA at an effective concentration of 80uM. (Note: If using an incubator, it is advisable to increase this incubation time to 40 minutes to account for the slower temperature change convection vs. conduction).
64. Near the end of the guide RNA incubation, thaw sufficient Cas9 protein to add 1:1 by volume to each well (10uL/well in this example) and warm to RT.
65. **Critical Step**: Using a multichannel pipettor, add 10uL Cas9 to the guide RNA mixture by moving the pipette tip in small, rapid circles while ejecting VERY SLOWLY. The 2:1 molar ratio of guide RNA to Cas9 promotes efficient crRNP formation, yielding a preparation of crRNPs at 20uM. **? Troubleshooting**: Rapid addition of the Cas9 may cause precipitation. If a precipitant forms, gently pipet up and down until the protein returns to suspension.
66. Reseal the plates and incubate at 37°C for 15 minutes to form crRNPs.
67. Using a multichannel pipettor, aliquot 3.5uL of crRNPs into 5 separate Lo-Bind 96-well V-bottom plates. Seal plates with foil, label, and store at −80°C until needed. **Pause Point**: crRNPs may be used immediately, but preparing crRNPs ahead of time dramatically improves screening workflow. Plates of crRNPs may be stored at −80°C in this buffer for at least 6 months or at 4°C for up to two weeks with little-to-no loss in editing efficiency. We strongly recommend that all crRNPs used in a screen be treated consistently (*i.e*. either all frozen at −80°C or all used immediately). Freeze/thaw of crRNPs is not advisable but limited cycles do not significantly impact editing efficiency. **CRISPR-Cas9 RNP Nucleofection Protocol**, Timing 1d **Timing**: If isolations are performed on Monday or Tuesday, nucleofections can be performed on Thursday or Friday of the same week allowing 72 hours for T cell activation.
68. Prepare the stimulatory beads from the T Cell Activation and Expansion Kit according to the manufacturer’s instructions. Allow the beads to rotate at 4°C at least 2 hours prior to use.
69. Open the P3 Primary Cell 4D-Nucleofector^®^ X Kit and mix the supplement and nucleofection buffer according to the manufacturers instructions. Supplemented P3 buffer can be stored at 4°C for up to 3 months. P3 buffer should be allowed to come to room temperature before it comes in contact with cells.
70. Set up the Amaxa 4D Nucleofection System according to manufacturer’s instructions. Use pulse code EH-115 for nucleofection.
71. Count the stimulated primary CD4+ T cells. Pool enough cells for 500,000 cells per nucleofection reaction and pellet at 400xg for 5 minutes in a spinning bucket centrifuge. For example, pellet 2 million cells for 4 reactions. **? Troubleshooting**: The editing protocol used here is effective for cell counts between 200,000 and 1 million cells per reaction. Cell number should be optimized for downstream application.
72. Thaw crRNPs in plate format or, if crRNPs are not in plate format, array 3.5uL (70pmol) of each crRNP in a 96-well Lo-Bind V-Bottom plate in the exact layout to be used for nucleofection. **Critical Step**: In order to minimize the amount of time the cells spend in nucleofection buffer, collect and arrange the necessary reagents beforehand for efficient handling.
73. Remove the supernatant from the pelleted cells and resuspend in 20uL nucleofection buffer per reaction. If performing many reactions, transfer cells to an appropriate vessel for multichannel pipetting.
74. Add 20uL cell suspension to the 3.5uL crRNPs and mix gently 3-4 times by pipet.
75. Transfer 20uL of each reaction to the nucleofection plate, leaving any extra solution behind. **? Troubleshooting**: To avoid arc errors during nucleofection, dispense the reaction at the bottom of the cuvettes and avoid air bubbles. Arc errors will not preclude successful nucleofection, but may negatively impact editing efficiency and/or cell viability.
76. Tap the nucleofection plate gently on the surface of the hood to release any bubbles that may have formed.
77. Nucleofect the cells on the Amaxa 4D-Nucleofector System using program EH-115.
78. As quickly as possible post-nucleofection, add 80uL fresh, pre-warmed cRPMI plus 20U/mL IL-2 to each well and allow the cells to recover for 15 minutes to 6 hours in the incubator while still in the nucleofection cuvette.
79. Meanwhile, prepare a flat-bottom 96-well plate with 100uL cRPMI per well plus 20U/mL IL-2 plus stimulatory beads at a 1:1 bead:cell ratio. Store in the incubator to keep warm until needed.
80. After the cells have been allowed to recover for at least 30 minutes in the nucleofection plate, transfer the entirety of each reaction from the nucleofection cuvette to the appropriate wells of the prepared, pre-warmed flat bottom plate for expansion in culture. **Cell Expansion and Validation**, Timing 4-8d **Timing**: To allow time for cell recovery, gene editing, and protein turnover, at least 3 days should pass prior to harvesting samples for validation. **Critical Step**: Edited cells will continue to expand for roughly 10-14 days post-treatment with stimulatory beads after which time the cells will lose activation and permissivity to infection. Cells can be expanded further with a second round of stimulation if necessary. The cells in this case are divided into 5 sets: one for harvesting protein, one for harvesting genomic DNA, and three for infection in technical triplicate.
81. 48-72 hours post-nucleofection, feed the cells by adding 100uL cRPMI plus 20U/mL IL-2 to each well of the plate.
82. 48-72 hours later, mix a representative well of cells by pipet and count cell density using a hemacytometer or cell counter. If 500,000 cells were nucleofected originally, expect the cells to be around 1-2 million cells/mL.
83. Using a multichannel pipet, suspend each well and transfer 100,000 cells each into two U-bottom, 96-well plates. Add back an equivalent volume of cRPMI + 20U/mL IL-2 to each well of the original plate and return these cells to the incubator.
84. Spin the U-bottom plates at 400xg for 3 minutes to pellet the cells. Gently remove the supernatant from each well.
85. Suspend cells in one plate in 60uL 2.5x Laemmli sample buffer per well. Transfer the lysates into PCR tubes and heat at 98°C for 20 minutes. These lysates can be used for immunoblot analysis to determine knock-out efficiency at the protein level (as in **Figure 4A**). Store lysates at −20°C.
86. Suspend cells in the second plate in 50uL QuickExtract buffer per well. Transfer the lysates into PCR tubes and heat at 65°C for 20 minutes followed by 98°C for 5 minutes. These lysates contain PCR-ready genomic DNA for editing analysis (see ‘TIDE Analysis’ below). Store lysates at −20°C. **Pause Point**: Edited primary CD4+ T cells can be frozen and stored in FBS +10% DMSO. Otherwise, they can be taken forward in phenotypic assays such as HIV spreading infections.
87. If proceeding with phenotypic analysis directly, cells can be plated immediately for HIV infection. Suspend a representative well of cells and count cell density using a hemacytometer or cell counter.
88. Using a multichannel pipet, resuspend each well and transfer at least 50,000 to 100,000 cells per well into each of three U-bottom, 96-well plates. These will be technical replicates for the infection. Plate a few extra wells of cells for un-infected controls.
89. Add enough cRMPI + 20U/mL IL-2 to each well in the U-bottom plates for a total volume of 150uL per well. Any extra cells can be used for additional validation, discarded, expanded, or cryopreserved.
90. **Critical Step**: It is advised to validate successful knock-out of a representative positive control prior to phenotypic analysis through immunostaining, immunoblotting, or TIDE (see below). In the case of CXCR4, extra non-targeting and *CXCR4*-targeted cells can be stained with CXCR4-APC in accordance to the manufacturer’s protocol and analyzed by flow cytometry (as in **Figure 3B**). Inefficient editing of the positive control likely indicates a recalcitrant donor or faulty crRNP synthesis. **TIDE (Tracking of Indels by Decomposition) Analysis**, Timing 3-4d
91. TIDE works by deconvoluting the Sanger sequencing chromatograms of polyclonal pools of DNA. CRISPR-Cas9 editing will result in the formation of randomly assorted insertions and deletions (Indels) at the cut site^37^. PCR amplification over the cut site from the genomic DNA of a polyclonal pool of cells provides a template for Sanger sequencing, which can then be analyzed for Indel percentage by the TIDE webtool^63^ (**Figure 3C**).
92. Design single-stranded DNA oligos to serve as PCR primers over the targeted cut site. Aim for a product size between 500 and 700 base pairs, with the guide RNA target site at least 150bp from either primer. Reference genome sequences for primer design can be found using UCSC genome browser (http://genome.ucsc.edu) or Ensembl (http://www.ensembl.org) and primers designed using a variety of webtools such as Primer3^64,65^ (http://bioinfo.ut.ee/primer3-0.4.0/primer3/) or Benchling (https://benchling.com/tutorials/25/creating-and-analyzing-primers). Each guide RNA will require unique TIDE primer pairs unless the target sites are in close proximity.
93. Suspend oligos in nuclease-free water and generate 10uM working stocks. Similarly, prepare working stocks of 10mM dNTPs.
94. Prepare 50uL PCR reactions using a high-fidelity polymerase (such as Phusion). For each primer pair, include amplification off the non-targeting control genomic DNA.

95. Amplify over the cut site using a touch-down PCR amplification strategy with appropriate annealing temperatures for the designed primers. For example: 98°C (5 minutes), [98°C (30 seconds), 65°C (20 seconds), 72°C (1 minute)] for 14 cycles with 0.5°C decrease in annealing temperature per cycle, [98°C (30 seconds), 58°C (20 seconds), 72°C (1 minute)] for 20 cycles, 72°C (10 minutes), 10°C (Final hold).
96. Purify the amplified DNA for sequencing using a PCR Purification Kit (*i.e*. Macherey Nagel’s Nucleospin PCR Clean-Up Kit) according to the manufacturer’s instructions.
97. Submit the PCR products for Sanger sequencing using both the forward and reverse TIDE oligos originally used for amplification.
98. Once the chromatograms are returned, sequence diversification will become evident directly adjacent to the PAM sequence in the guide if the nucleofection was successful (*i.e*. **Figure 3C**).
99. An estimated percentage of Indels can be generated by uploading the experimental and control chromatograms to the TIDE webtool online (http://tide.nki.nl). The best guides will reveal allelic editing percentages greater than 75% by TIDE. **Caution**: TIDE data should not be considered absolute empirical knock-out percentages, but instead provides a rough estimate of editing for rapid validation of editing. **Generation of Concentrated HIV-1 Virus Stocks**, Timing 5d **Timing**: To allow time for virus generation, precipitation, and titering, viruses should be made no later than the week of primary cell isolation, preferably earlier. Once precipitated and stored, the viruses will be stable at −80°C for several months. Virus generation takes 5 days and so Monday is the recommended start day (**Figure 2**).

| <u>Reagent</u> | <u>uL</u> |
| --- | --- |
| 5X Buffer | 10 |
| dNTPs (10mM) | 2 |
| dH2O | 31 |
| F primer (10uM) | 2 |
| R primer (10uM) | 2 |
| Phusion | 0.5 |
| Genomic DNA | 2.5 |
|  | 50uL total |

**Figure 2.**
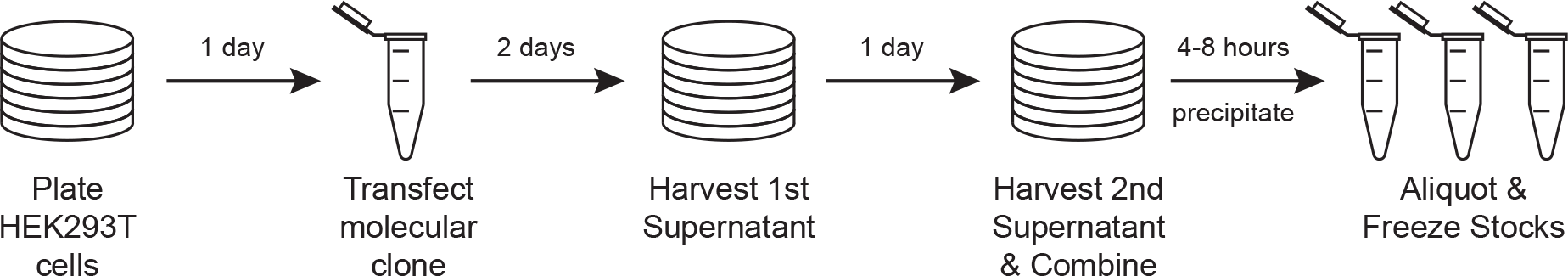
Experimental overview of HIV-1 virus preparation. HEK293T cells are plated and transfected with an HIV-1 molecular clone using appropriate biosafety precautions. Virus containing supernatant is harvested 48 and 72 hours after transfection. Polyethylene glycol precipitation followed by centrifugation is used to precipitate the virus for concentrated resuspension, titering, and storage.
100. Starting with a confluent T175 flask of HEK293T cells, remove the media, rinse gently with 10mL 1xPBS, and add 5mL 0.05% trypsin. Gently rock for 2-3 minutes until the cells lift off the flask.
101. Pellet the cells in a 50mL conical in a spinning bucket rotor at 400xg for 3 minutes. Remove the supernatant and suspend the cells in 50mL DMEM media.
102. Calculate cell density using a hemacytometer or cell counter. Transfer 5 million cells into each of eight 15cm plates and add prewarmed DMEM media to a final volume of 27mL. Allow cells to settle and adhere overnight.
103. The following day, prepare 8 microcentrifuge tubes, one per 15cm dish, with 10ug replication-competent proviral vector in accordance with the experimental design. This protocol uses an HIV-1 NL4-3 full molecular clone with an integrated GFP reporter driven off of an IRES sequence following the *nef* open reading frame.
104. Warm serum-free DMEM and polyJET to room temperature for transfection.
105. Add 250uL serum-free DMEM (25uL per ug DNA) to each DNA tube.
106. Prepare a master mix of 250uL serum-free DMEM (25uL per ug DNA) and 30uL polyJET reagent (3uL per ug DNA) per tube. For eight plates, this is 2mL serum-free DMEM + 240uL polyJet. Incubate for 2 minutes at RT.
107. Add 250uL polyJET mixture to each diluted DNA mixture and pipet to mix. Incubate at RT for 15 minutes.
108. Add each transfection mixture dropwise to cells. Swirl dish to mix and return the cells to the incubator. **Caution**: As soon as the cells are transfected, they are considered HIV-positive and therefore biohazardous and must be handled according to the appropriate biosafety standards at the respective institution.
109. After 48 hours, remove the 27mL viral supernatant from each plate and store in 50mL conicals at 4°C. Replace immediately with 25mL fresh complete DMEM media and return the cells to the incubator overnight.
110. The following day, remove the second batch of viral supernatant and combine with the first. Filter out cell debris from the viral supernatant through a PVDF filter (i.e. Steriflip device or similar). Bleach the virus producing cells and dispose.
111. Meanwhile, prepare 16 50mL conicals for virus precipitation. To each tube, add 5.5mL 50% PEG-6000 (to achieve 8.5% final concentration) and 2.4mL 4M NaCl (to achieve 0.3M final concentration) for virus precipitation. **Timing**: PEG-6000 and NaCl solution may be aliquoted the night before.
112. To each tube, add 24mL filtered virus supernatant. Invert to mix and let virions precipitate at 4°C for at least 4 hours but not more than 8 hours.
113. Pellet the virions by centrifugation in a spinning bucket rotor at 3500rpm for 20 minutes at 4°C.
114. Immediately after spinning, decant or aspirate off the supernatant and add 250uL PBS to the precipitate in each tube. **Caution**: Avoid aerosols by spinning in secondary containment and opening the centrifuge buckets only in a biosafety cabinet. Decontaminate the inside of the secondary containment thoroughly before returning it to the centrifuge.
115. Pipet to resuspend and combine all fractions. Expected yield will be roughly 6mL virus stock at 100-fold concentration. **Caution**: Concentrated, replication-competent HIV-1 stocks must be handled with extreme care according to appropriate biosafety standards at the respective institution.
116. Aliquot the virus into approved 1.5mL screw-top, microcentrifuge tubes for storage. Plan for roughly 24-36 aliquots with 50, 100, or 250uL of virus. Snap freeze on dry ice for best yield. **Pause Point**: Virus can be concentrated and stored at −80°C for at least 6 months without a loss of viability.
117. A sample of frozen virus can be thawed for virus titer calculation. **Critical Step**: Due to donor variability in virus susceptibility (i.e. see Anticipated Results, **Figure 4B**), it is recommended that viral stocks be quantified as p24 equivalents or by donor-matched live titer. Live titers can be estimated by infecting stimulated primary T cells over a dilution series of virus while p24 levels in the viral stock can be quantified using a variety of commercially available kits (*i.e*. Cell Biolab’s QuickTiter™ HIV Lentivirus Quantitation Kit). **Primary Cell HIV Spreading Infection**, Timing 7-9d **Timing**: While spreading infection timepoints may be taken with some flexibility, it is generally advantageous to take at least three between 3-7 days post-infection. If initial infection occurs on Friday, this permits sampling on Monday, Wednesday, and Friday of the following week. Three timepoints in technical triplicate will ultimately result in nine flow samples per nucleofected population. **Critical Step**: To ensure successful infection, the cells must be activated at the time of virus exposure. Therefore stimulation should occur at least 3-7 days prior to infection. **? Troubleshooting**: If infection rates are too low, spinoculation of cultures can be a viable alternative to spreading infection. Full molecular clones without integrated reporters can also be analyzed by p24 staining and flow cytometry or by monitoring of p24 in the culture supernatant by ELISA (*i.e*., ^66^).
118. Based on titer results after viral stock generation, calculate the required amount of virus to add to each well to achieve roughly a 1-2% initial infection after 72 hours. For most donors and viruses, this will be roughly 2-5uL of concentrated virus per well.
119. In a 50mL conical, aliquot sufficient cRPMI plus 20U/mL IL-2 to add 50uL per well of cells. Thaw and add the appropriate amount of virus stock to this tube. Recall that cells were plated in technical triplicate over three identical plates.
120. Mix the virus and media gently and add 50uL of virus inoculate to each well across each plate. Don’t forget to add 50uL RPMI media without virus to the extra uninfected control wells. Return to the incubator for 72 hours.
121. Prior to taking the first time point, prepare 75uL per well of 2% formaldehyde in PBS for cell fixation. Label 96-well U-bottom plates for fixation and aliquot 75uL of 2% formaldehyde per well. Additionally, prepare enough cRPMI plus 20U/mL IL-2 to add 75uL per well.
122. Gently mix each infected well with a multichannel pipet and remove 75uL mixed culture to the prepared fixation plate for a final concentration of 1% formaldehyde. Pipette up and down in the formaldehyde solution to prevent clumping.
123. Add 75uL fresh cRPMI plus 20U/mL IL-2 to each well of the infected plate and return to the incubator for 48 hours.
124. Wait 30 minutes for complete fixation and inactivation of virus. Clean the outside of the plate with 70% ethanol and wrap it in aluminum foil. Fixed cells can be removed for flow cytometry, gating both on live cells and GFP+ (infected) cells. **Pause Point**: Fixed cells can be stored in foil at 4°C for up to one month prior to flow cytometry.
125. Repeat steps 121-124 for the second timepoint (typically 5 days post-infection).
126. For the third and final timepoint, prepare 50uL per well of 4% formaldehyde in PBS for cell fixation. Label 96-well U-bottom plates for fixation and aliquot 50uL of 4% formaldehyde per well.
127. Gently mix each well with a multichannel pipet and remove 150uL mixed culture to the prepared fixation plate for a final concentration of 1% formaldehyde. Pipette up and down in the formaldehyde solution to prevent clumping.
128. Remaining cultures can be disposed of after inactivation with 10% bleach for 30 minutes.
129. Wait 30 minutes for complete fixation and inactivation of virus. Clean the outside of the fixation plate with 70% ethanol and wrap it in aluminum foil. Fixed cells can be removed for flow cytometry.
130. Analyze results by flow cytometry and appropriate analysis software (*i.e*. FlowJo), calculating percent of live cells and percent of infected cells over the timecourse of infection (**Figure 4B**).

## TIMING

Steps 1-11, Designing CRISPR guides: 1d
Steps 12-17, Resuspending CRISPR guides: 1d
Steps 18-33, PBMC Isolation: 1d
Steps 34-42, CD4+ T cell enrichment: 1d
Steps 43-49, T cell activation: 2-3d
Steps 50-56, Generating CRISPR-Cas9 RNPs, Small Scale Tubes: 1d
Steps 57-67, Generating CRISPR-Cas9 RNPs, Large Scale Plates: 1d
Steps 68-80, Nucleofection: 1d
Steps 81-90, Cell expansion and validation: 4-8d
Steps 91-99, TIDE analysis: 3-4d
Steps 100-117, Generating concentrated HIV-1 stocks: 5d
Steps 118-130, HIV spreading infection: 7-9d

## ANTICIPATED RESULTS

The results from these experiments will vary widely dependent on the biological question, donor sample, and phenotypic output. Following the protocol as outlined above, three critical pieces of data will be generated: quality control staining for CD4+ T cell isolation and stimulation, molecular validation of gene knock-out (by immunostaining, immunoblotting, or TIDE), and HIV spreading infection profiles in the edited T cell populations.

### CD4+ T Cell Isolation and Stimulation

Three primary cell samples are collected during isolation and stimulation at distinct stages: after Ficoll centrifugation (PBMCs, Step 36), after CD4+ T cell isolation (CD4+ enriched, Step 46), and 72 hours after stimulation (post-stimulation, Step 52). These are then stained for CD4 and CD25 to ensure proper isolation and activation, respectively (Step 53, **Figure 3A**). PBMCs from the whole blood of a typical donor under normal conditions will contain around 525% CD4+ T cells, only a fraction of which will be stimulated (5-20% CD25+). Most enrichment techniques are sufficient to purify CD4+ T cells to >95% of the population. Negative selection techniques for the isolation of naïve T cells will return only unstimulated (<5% CD25+) cells, whereas positive selection techniques will yield a ratio of stimulated:unstimulated cells similar to what was observed in the PBMC fraction. Post-stimulation, all isolated cells retain CD4 expression and may even increase in signal due to the larger cell size and surface area. Most protocols for stimulation (such as treatment with anti-CD3/anti-CD28/IL-2 as above) will yield near complete activation (>90% CD25+). As the editing of most loci is dependent on T cell activation, ensuring high levels of stimulation prior to nucleofection is critical to ensure success of the downstream protocols.

**Figure 3.**
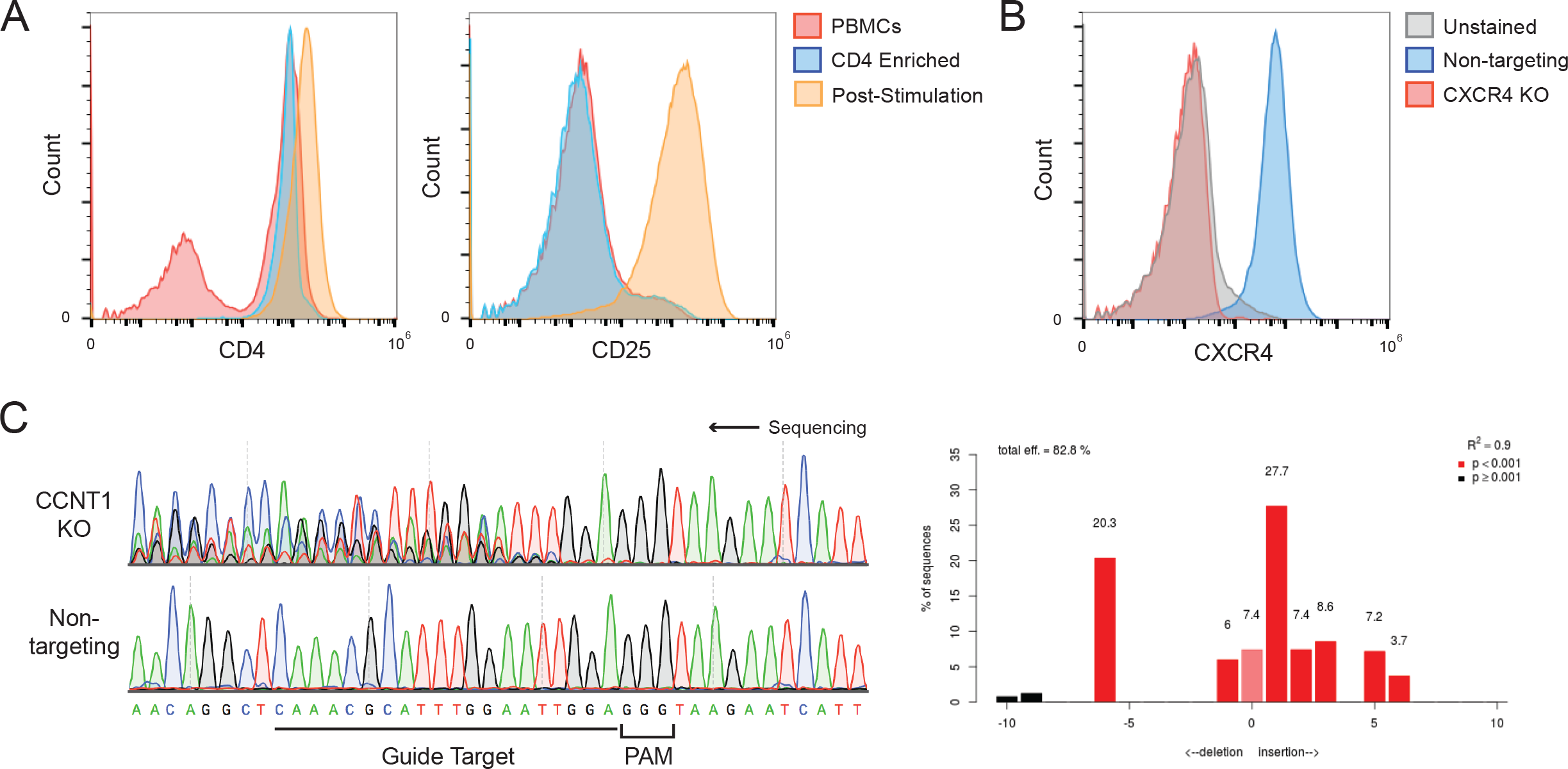
Anticipated results from primary T cell isolation and editing. **(a)** Flow cytometry histograms depicting CD4 and CD25 levels on the cell surface of PBMCs (red), CD4+ T cells post-enrichment (blue), and CD4+ T cells post-stimulation with anti-CD3/anti-CD28/IL-2 (orange). **(b)** Flow cytometry histogram depicting CXCR4 levels on the cell surface of primary T cells treated with non-targeting crRNPs (blue) or CXCR4-targeting crRNPs (red) relative to an unstained control (gray). **(c)** Chromatograms from Sanger sequencing of PCR amplicons over a target site in the CCNT1 locus in non-targeting treated cells (bottom) and CCNT1 crRNP treated cells (top). The guide RNA target sequence and associated PAM are highlighted below. The TIDE output calculating percent Indels from these chromatograms is depicted on the right.

### Validation of Gene Editing

A variety of molecular biology techniques can be employed to verify successful knockout of target genes including DNA sequencing, immunoblotting, and immunostaining. While edits at the DNA level can be observed within 24 hours of nucleofection, we recommend waiting at least 3 to 4 days post-nucleofection for validation. This not only ensures that editing has been completed, but also allows time for cell recovery post-nucleofection, the resolution of persistent DNA breaks, and the turnover of protein pools.

For large numbers of samples with unique molecular readouts, it may be advantageous to complete phenotypic analysis prior to validation of each condition depending on the experimental setup. Regardless of the design, we strongly recommend validation of the CXCR4 positive control prior to proceeding. The *CXCR4* locus has proven to be highly susceptible to editing in primary T cells and serves as a surrogate marker for efficiency. Relative to nontargeting controls, we expect ablation of CXCR4 from the cell surface of almost all cells as monitored by immunostaining (**Figure 3B**). Inefficient knock-out at the *CXCR4* locus likely indicates either an experimental error or a donor sample less susceptible to editing.

For proteins that lack appropriate antibodies for immunostaining or immunoblotting, DNA sequencing of the targeted site provides a good indication of successful editing. As the break sites within the polyclonal pool are unique to each locus and cell, PCR amplification over the targeted region will result in many unique fragments of differing length and sequence. These fragments can be precisely quantified by a number of deep sequencing approaches, but the easiest way to quickly determine rough percent editing is through Tracking of Indels by Decomposition (TIDE, Steps 95-103). This approach uses computational methods to deconvolute Sanger sequencing of related, yet diverse, PCR amplicons. For example, **Figure 3C** (right) shows the Sanger sequencing results from PCR amplicons over a targeted region of the *CCNT1* locus in both experimental (top) and non-targeting control (bottom) samples. All amplicons in the non-targeting control return the same sequence. In the experimental sample, however, the sequences diverge at the targeted cut site, indicating successful editing and the presence of multiple insertions and deletions (Indels). Uploading these chromatograms into the TIDE software returns an estimated total efficiency, including predicted frame shift percentages (**Figure 3C**, right). TIDE analysis is designed solely to give an estimate of Indel percentage and does not take into account nonsense mutations and other aberrant sequences. Nevertheless, it is a quick and cost effective method to validate site specific editing.

Editing efficiency will vary by locus, guide RNA, cell type, and donor, so it is vital to validate knock-out efficiency in every unique sample. Due to the variability inherent in working with these primary cell types, it is critical to validate any pertinent results with multiple guide RNA in multiple donors.

### HIV Spreading Infection

While many approaches have been developed for monitoring HIV replication in *ex vivo* cultures (p24 ELISA, reverse transcription assays, luciferase reporter viruses, fluorescent p24 antibodies, etc), we use a Nef:IRES:GFP reporter virus for phenotypic readout for three reasons. First, this virus is replication-competent and thus we can capture defects that occur both early and late in the infection cycle. Second, this strain expresses all viral open reading frames alongside an easily quantifiable marker without additional sample processing. Third, the use of flow cytometry to detect GFP provides single cell data (both percent GFP positive as well as mean fluorescent intensity), which can inform downstream mechanistic assays. The collection of single cell data also allows for the simultaneous monitoring of cell viability, shape, and size such that potential confounding effects in gene editing may be readily identified. Inclusion of a toxicity control, such as a *CDK9* targeting guide, can assist in the identification of similarly essential genes.

We recommend taking at least three timepoints post-infection, including one within the first 2 to 3 days. This first timepoint will ideally occur before the virus has completed multiple rounds of replication and is a surrogate for initial infection rate. Ideally, infection at day 2 or 3 will be between 1% and 3%, low enough that the virus can spread effectively without inducing massive cytotoxic effects in the culture. As the virus replicates, cell death and the relaxation of the cells in culture over time will eventually decrease percent infected cells, with peak infectivity typically occurring between days 5 and 9 post-infection.

Even prior to editing, each donor will be differentially susceptible to infection and baseline infection rates will vary. As such, it is essential to interpret each assay relative to donor-matched controls. For example, the results of an HIV spreading infection assay in two different donors are shown in **Figure 4E**. The baseline infection rate in the non-targeting control is roughly twofold lower for Donor B than for Donor A though they were treated identically and in parallel. Nevertheless, the results are consistent between the two donors. As expected, knockout of the *CXCR4* (dark blue triangle) and *LEDGF* (light blue triangle) controls dramatically repressed replication relative to the non-targeting control (gray square). Three experimental guides that target *CCNT1* were tried in each donor (red circles). These guides appear differentially effective in inhibiting HIV replication, with one guide (dark red, #3) inhibiting replication to near control levels. Immunoblots probed for CCNT1 protein levels in these samples (**Figure 4D**) confirm that the most effective guide gave the strongest HIV replication phenotype.

**Figure 4.**
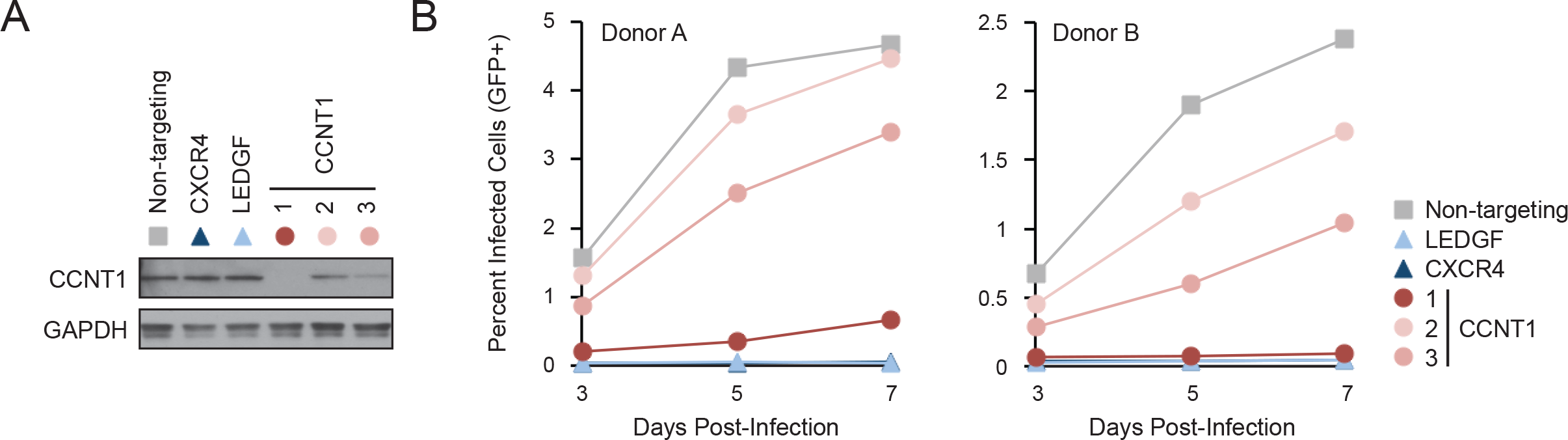
Anticipated results from primary T cell spreading infection. **(a)** An immunoblot of CCNT1 levels in primary T cells treated with 6 different crRNPs: non-targeting (gray square), CXCR4 (dark blue triangle), LEDGF (light blue triangle), and three unique guides targeting CCNT1 (red circles); relative to a GAPDH loading control. **(b)** Average percent HIV infected (GFP+) cells at days 3, 5, and 7 post-infection are shown for 6 pools of edited cells in two donors (n=3 technical triplicates).

Correlating guide strength to phenotype strength and confirming the phenotype in multiple donors is critical to interpretation of the results. For screens with large numbers of samples, we recommend performing these assays in at least two donors and normalizing the infectivity relative to the median infectivity across the plate at a given timepoint for a given donor. Assuming most guides will not impact replication, the distribution about the median will allow for the determination of statistical cut offs. If a large number of guides are anticipated to have an impact, normalization relative to the average of the non-targeting controls is another option. Critical findings should be repeated in multiple donors after the screening is completed with consideration for phenotypic variability, strength, and statistical robustness.

## ACKNOWLEDGEMENTS

We thank Lars Pache, Emilie Battivelli, and members of the Marson and Krogan labs for critical feedback and testing of the protocol. This research was supported by amfAR grant 109504-61-RKRL using funds raised by generationCURE (J.F.H.), a fellowship of the Deutsche Forschungsgemeinschaft (SCHU3020/2-1; K.S.), the UCSF Sandler Fellowship (A.M.), a gift from Jake Aronov (A.M.), NIH/NIGMS funding for the HIV Accessory & Regulatory Complexes (HARC) Center (P50 GM082250, A.M., J.D. and N.J.K.), NIH funding for the FluOMICs cooperative agreement (U19 AI106754, J.F.H. and N.J.K.), NIH/NIAID funding for the HIV Immune Networks Team (P01 AI090935, N.J.K.), NIH funding for the Dengue Human Immunology Project Consortium (DHIPC, U19 AI118610, N.J.K.), NIH funding for the study of innate immune responses to intracellular pathogens (R01 AI120694 & P01 AI063302, N.J.K.), NIH funding for the UCSF-Gladstone Institute of Virology & Immunology Center for AIDS Research (CFAR, P30 AI027763), and an NIH/NIDA grant (DP2 DA042423-01, A.M.). A.M. holds a Career Award for Medical Scientists from the Burroughs Wellcome Fund and is a Chan Zuckerberg Biohub Investigator. Special thanks to Ethan Brookes, Matthew Hall, and Olivier Cantada at Lonza Bioscience for their support with the nucleofection transfection technology, Darrick Chow at Dharmacon for his support with guide RNA synthesis, and Chris Jeans at the University of California, Berkeley Macrolab for the production of Cas9 protein.

## AUTHOR CONTRIBUTIONS

J.F.H, J.H., K.S., N.J.K, and A.M designed the experimental procedure, which was further optimized with input from J.H., T.L.R., M.J.M., P.H., and J.D. Quality control and infection data were collected by J.F.H., J.H., and M.J.M. The figures were designed and assembled by J.F.H. with input from J.H. Text was written by J.F.H., J.H., A.M., and N.J.K. with critical input from M.J.M., P.H., K.S., T.L.R., and J.D. Correspondence and requests for materials should be addressed to.

## ? TROUBLESHOOTING

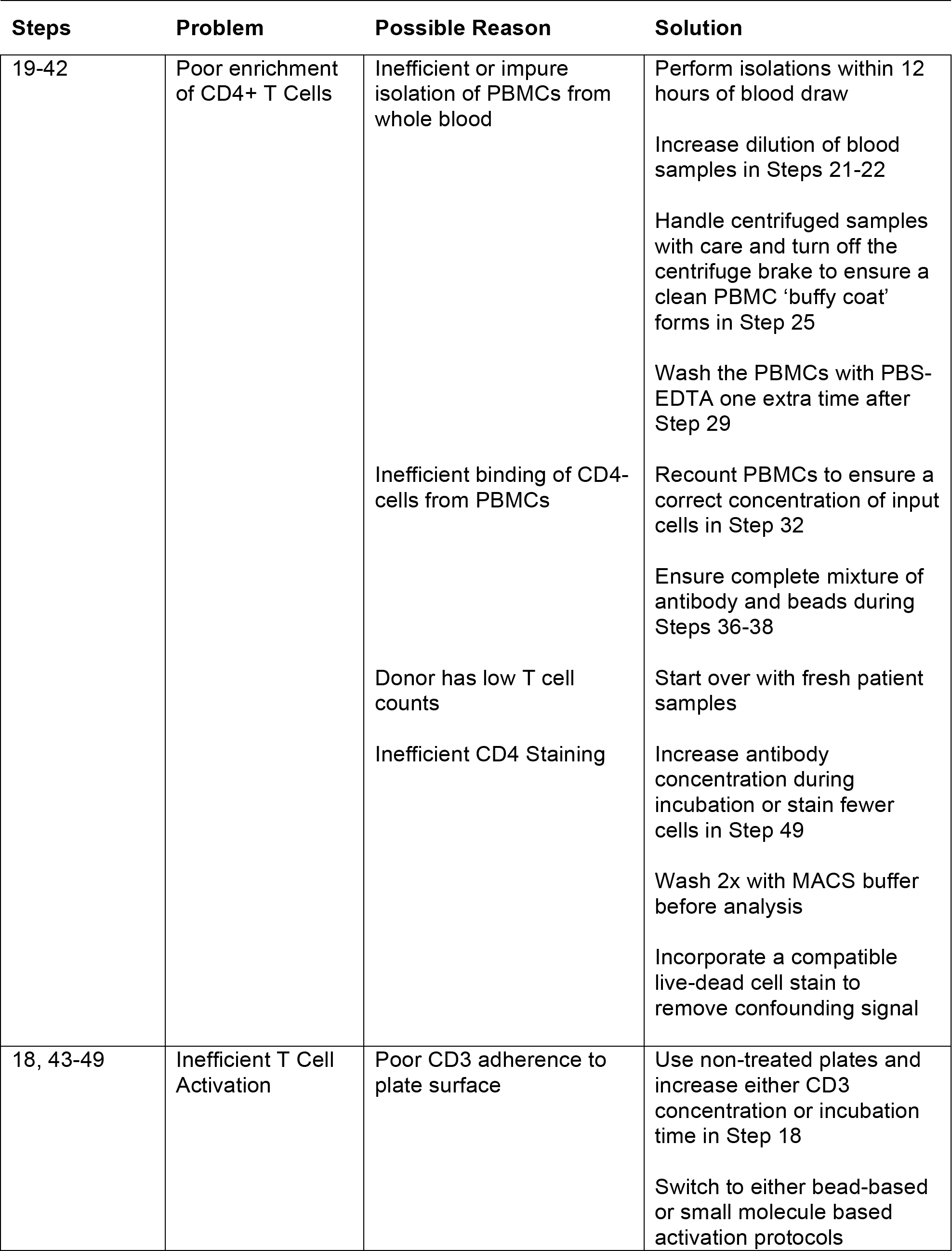
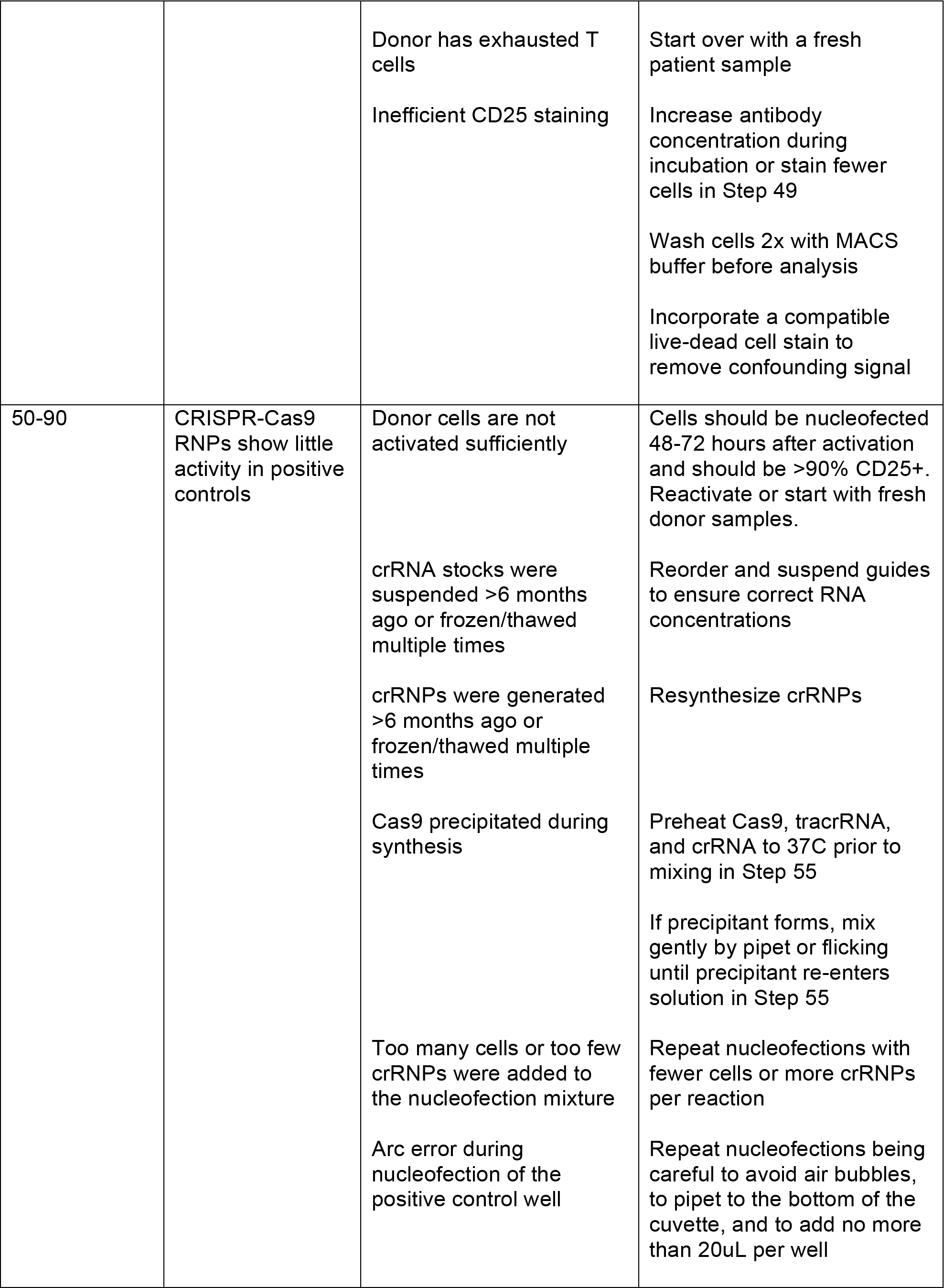
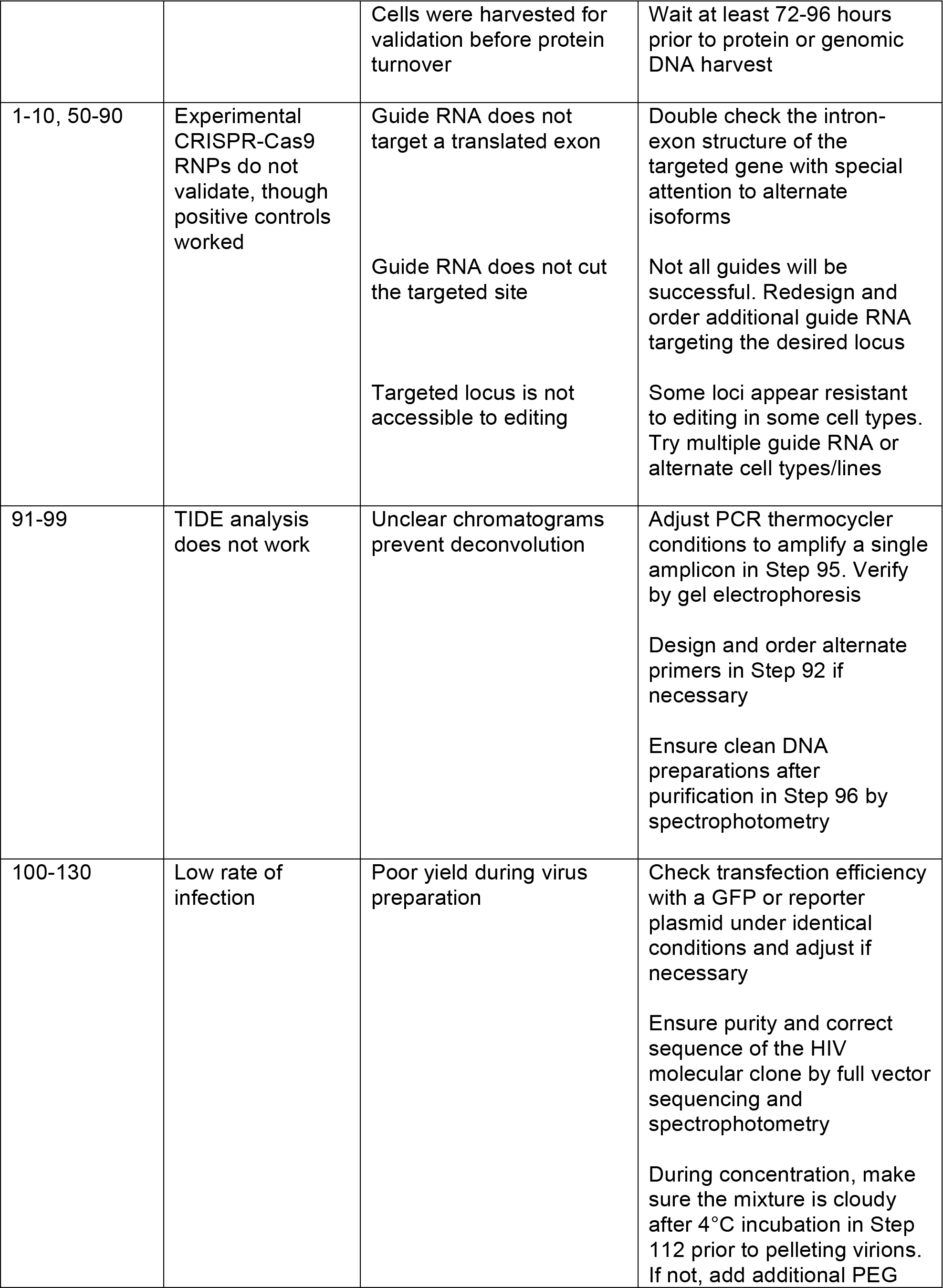
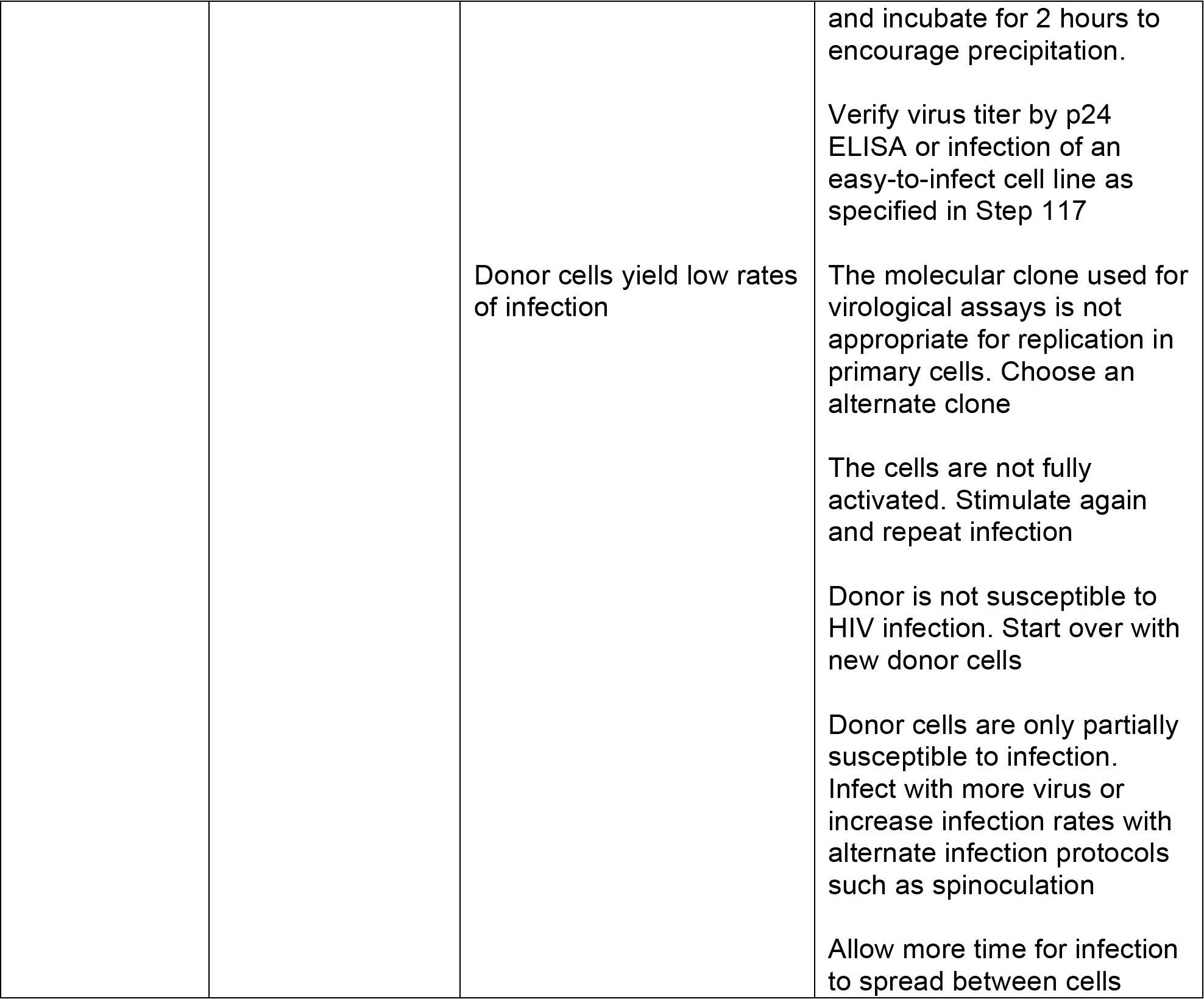

## REFERENCES

1 Harris, R.S., Hultquist, J.F., & Evans, D.T., The restriction factors of human immunodeficiency virus. The Journal of biological chemistry 287 (49), 40875–40883 (2012).

2 Goff, S.P., Host factors exploited by retroviruses. Nature reviews. Microbiology 5 (4), 253–263 (2007).

3 Hsu, T.H. & Spindler, K.R., Identifying host factors that regulate viral infection. PLoS pathogens 8 (7), e1002772 (2012).

4 Louz, D., Bergmans, H.E., Loos, B.P., & Hoeben, R.C., Animal models in virus research: their utility and limitations. Critical reviews in microbiology 39 (4), 325–361 (2013).

5 Ambrose, Z., KewalRamani, V.N., Bieniasz, P.D., & Hatziioannou, T., HIV/AIDS: in search of an animal model. Trends in biotechnology 25 (8), 333–337 (2007).

6 Hatziioannou, T. & Evans, D.T., Animal models for HIV/AIDS research. Nature reviews. Microbiology 10 (12), 852–867 (2012).

7 Bushman, F.D., Malani, N., Fernandes, J., D'Orso, I., Cagney, G., Diamond, T.L., Zhou, H., Hazuda, D.J., Espeseth, A.S., Konig, R., Bandyopadhyay, S., Ideker, T., Goff, S.P., Krogan, N.J., Frankel, A.D., Young, J.A., & Chanda, S.K., Host cell factors in HIV replication: meta-analysis of genome-wide studies. PLoS pathogens 5 (5), e1000437 (2009).

8 Yeung, M.L., Houzet, L., Yedavalli, V.S., & Jeang, K.T., A genome-wide short hairpin RNA screening of jurkat T-cells for human proteins contributing to productive HIV-1 replication. The Journal of biological chemistry 284 (29), 19463–19473 (2009).

9 Park, R.J., Wang, T., Koundakjian, D., Hultquist, J.F., Lamothe-Molina, P., Monel, B., Schumann, K., Yu, H., Krupzcak, K.M., Garcia-Beltran, W., Piechocka-Trocha, A., Krogan, N.J., Marson, A., Sabatini, D.M., Lander, E.S., Hacohen, N., & Walker, B.D., A genome-wide CRISPR screen identifies a restricted set of HIV host dependency factors. Nature genetics 49 (2), 193–203 (2017).

10 Lorsch, J.R., Collins, F.S., & Lippincott-Schwartz, J., Cell Biology. Fixing problems with cell lines. Science 346 (6216), 1452–1453 (2014).

11 Hughes, P., Marshall, D., Reid, Y., Parkes, H., & Gelber, C., The costs of using unauthenticated, over-passaged cell lines: how much more data do we need? BioTechniques 43 (5), 575, 577–578, 581-572 passim (2007).

12 American Type Culture Collection Standards Development Organization Workgroup, A.S.N., Cell line misidentification: the beginning of the end. Nature reviews. Cancer 10 (6), 441–448 (2010).

13 Pan, C., Kumar, C., Bohl, S., Klingmueller, U., & Mann, M., Comparative proteomic phenotyping of cell lines and primary cells to assess preservation of cell type-specific functions. Molecular & cellular proteomics: MCP 8 (3), 443–450 (2009).

14 Alge, C.S., Hauck, S.M., Priglinger, S.G., Kampik, A., & Ueffing, M., Differential protein profiling of primary versus immortalized human RPE cells identifies expression patterns associated with cytoskeletal remodeling and cell survival. Journal of proteome research 5 (4), 862–878 (2006).

15 Leroy, B., Girard, L., Hollestelle, A., Minna, J.D., Gazdar, A.F., & Soussi, T., Analysis of TP53 mutation status in human cancer cell lines: a reassessment. Human mutation 35 (6), 756–765 (2014).

16 Hultquist, J.F., Schumann, K., Woo, J.M., Manganaro, L., McGregor, M.J., Doudna, J., Simon, V., Krogan, N.J., & Marson, A., A Cas9 Ribonucleoprotein Platform for Functional Genetic Studies of HIV-Host Interactions in Primary Human T Cells. Cell reports 17 (5), 1438–1452 (2016).

17 Schumann, K., Lin, S., Boyer, E., Simeonov, D.R., Subramaniam, M., Gate, R.E., Haliburton, G.E., Ye, C.J., Bluestone, J.A., Doudna, J.A., & Marson, A., Generation of knock-in primary human T cells using Cas9 ribonucleoproteins. Proceedings of the National Academy of Sciences of the United States of America 112 (33), 10437–10442 (2015).

18 Brass, A.L., Dykxhoorn, D.M., Benita, Y., Yan, N., Engelman, A., Xavier, R.J., Lieberman, J., & Elledge, S.J., Identification of host proteins required for HIV infection through a functional genomic screen. Science 319 (5865), 921–926 (2008).

19 Konig, R., Zhou, Y., Elleder, D., Diamond, T.L., Bonamy, G.M., Irelan, J.T., Chiang, C.Y., Tu, B.P., De Jesus, P.D., Lilley, C.E., Seidel, S., Opaluch, A.M., Caldwell, J.S., Weitzman, M.D., Kuhen, K.L., Bandyopadhyay, S., Ideker, T., Orth, A.P., Miraglia, L.J., Bushman, F.D., Young, J.A., & Chanda, S.K., Global analysis of host-pathogen interactions that regulate early-stage HIV-1 replication. Cell 135 (1), 49–60 (2008).

20 Zhou, H., Xu, M., Huang, Q., Gates, A.T., Zhang, X.D., Castle, J.C., Stec, E., Ferrer, M., Strulovici, B., Hazuda, D.J., & Espeseth, A.S., Genome-scale RNAi screen for host factors required for HIV replication. Cell host & microbe 4 (5), 495–504 (2008).

21 Pache, L., Konig, R., & Chanda, S.K., Identifying HIV-1 host cell factors by genome-scale RNAi screening. Methods 53 (1), 3–12 (2011).

22 Llano, M., Saenz, D.T., Meehan, A., Wongthida, P., Peretz, M., Walker, W.H., Teo, W., & Poeschla, E.M., An essential role for LEDGF/p75 in HIV integration. Science 314 (5798), 461–464 (2006).

23 Llano, M., Vanegas, M., Fregoso, O., Saenz, D., Chung, S., Peretz, M., & Poeschla, E.M., LEDGF/p75 determines cellular trafficking of diverse lentiviral but not murine oncoretroviral integrase proteins and is a component of functional lentiviral preintegration complexes. Journal of virology 78 (17), 9524–9537 (2004).

24 Fadel, H.J., Morrison, J.H., Saenz, D.T., Fuchs, J.R., Kvaratskhelia, M., Ekker, S.C., & Poeschla, E.M., TALEN knockout of the PSIP1 gene in human cells: analyses of HIV-1 replication and allosteric integrase inhibitor mechanism. Journal of virology 88 (17), 9704–9717 (2014).

25 Shun, M.C., Raghavendra, N.K., Vandegraaff, N., Daigle, J.E., Hughes, S., Kellam, P., Cherepanov, P., & Engelman, A., LEDGF/p75 functions downstream from preintegration complex formation to effect gene-specific HIV-1 integration. Genes & development 21 (14), 1767–1778 (2007).

26 Jackson, A.L. & Linsley, P.S., Recognizing and avoiding siRNA off-target effects for target identification and therapeutic application. Nature reviews. Drug discovery 9 (1), 57–67 (2010).

27 Jackson, A.L., Burchard, J., Schelter, J., Chau, B.N., Cleary, M., Lim, L., & Linsley, P.S., Widespread siRNA “off-target” transcript silencing mediated by seed region sequence complementarity. Rna 12 (7), 1179–1187 (2006).

28 Tripathi, S., Pohl, M.O., Zhou, Y., Rodriguez-Frandsen, A., Wang, G., Stein, D.A., Moulton, H.M., DeJesus, P., Che, J., Mulder, L.C., Yanguez, E., Andenmatten, D., Pache, L., Manicassamy, B., Albrecht, R.A., Gonzalez, M.G., Nguyen, Q., Brass, A., Elledge, S., White, M., Shapira, S., Hacohen, N., Karlas, A., Meyer, T.F., Shales, M., Gatorano, A., Johnson, J.R., Jang, G., Johnson, T., Verschueren, E., Sanders, D., Krogan, N., Shaw, M., Konig, R., Stertz, S., Garcia-Sastre, A., & Chanda, S.K., Meta-and Orthogonal Integration of Influenza “OMICs” Data Defines a Role for UBR4 in Virus Budding. Cell host & microbe 18 (6), 723–735 (2015).

29 Pichlmair, A., Kandasamy, K., Alvisi, G., Mulhern, O., Sacco, R., Habjan, M., Binder, M., Stefanovic, A., Eberle, C.A., Goncalves, A., Burckstummer, T., Muller, A.C., Fauster, A., Holze, C., Lindsten, K., Goodbourn, S., Kochs, G., Weber, F., Bartenschlager, R., Bowie, A.G., Bennett, K.L., Colinge, J., & Superti-Furga, G., Viral immune modulators perturb the human molecular network by common and unique strategies. Nature 487 (7408), 486–490 (2012).

30 Carette, J.E., Guimaraes, C.P., Varadarajan, M., Park, A.S., Wuethrich, I., Godarova, A., Kotecki, M., Cochran, B.H., Spooner, E., Ploegh, H.L., & Brummelkamp, T.R., Haploid genetic screens in human cells identify host factors used by pathogens. Science 326 (5957), 1231–1235 (2009).

31 Deans, R.M., Morgens, D.W., Okesli, A., Pillay, S., Horlbeck, M.A., Kampmann, M., Gilbert, L.A., Li, A., Mateo, R., Smith, M., Glenn, J.S., Carette, J.E., Khosla, C., & Bassik, M.C., Parallel shRNA and CRISPR-Cas9 screens enable antiviral drug target identification. Nature chemical biology 12 (5), 361–366 (2016).

32 Heaton, N.S., Moshkina, N., Fenouil, R., Gardner, T.J., Aguirre, S., Shah, P.S., Zhao, N., Manganaro, L., Hultquist, J.F., Noel, J., Sachs, D., Hamilton, J., Leon, P.E., Chawdury, A., Tripathi, S., Melegari, C., Campisi, L., Hai, R., Metreveli, G., Gamarnik, A.V., Garcia-Sastre, A., Greenbaum, B., Simon, V., Fernandez-Sesma, A., Krogan, N.J., Mulder, L.C.F., van Bakel, H., Tortorella, D., Taunton, J., Palese, P., & Marazzi, I., Targeting Viral Proteostasis Limits Influenza Virus, HIV, and Dengue Virus Infection. Immunity 44 (1), 46–58 (2016).

33 Doudna, J.A. & Charpentier, E., Genome editing. The new frontier of genome engineering with CRISPR-Cas9. Science 346 (6213), 1258096 (2014).

34 Hsu, P.D., Lander, E.S., & Zhang, F., Development and applications of CRISPR-Cas9 for genome engineering. Cell 157 (6), 1262–1278 (2014).

35 Ran, F.A., Hsu, P.D., Wright, J., Agarwala, V., Scott, D.A., & Zhang, F., Genome engineering using the CRISPR-Cas9 system. Nature protocols 8 (11), 2281–2308 (2013).

36 Mandal, P.K., Ferreira, L.M., Collins, R., Meissner, T.B., Boutwell, C.L., Friesen, M., Vrbanac, V., Garrison, B.S., Stortchevoi, A., Bryder, D., Musunuru, K., Brand, H., Tager, A.M., Allen, T.M., Talkowski, M.E., Rossi, D.J., & Cowan, C.A., Efficient ablation of genes in human hematopoietic stem and effector cells using CRISPR/Cas9. Cell stem cell 15 (5), 643–652 (2014).

37 van Overbeek, M., Capurso, D., Carter, M.M., Thompson, M.S., Frias, E., Russ, C., Reece-Hoyes, J.S., Nye, C., Gradia, S., Vidal, B., Zheng, J., Hoffman, G.R., Fuller, C.K., & May, A.P., DNA Repair Profiling Reveals Nonrandom Outcomes at Cas9-Mediated Breaks. Molecular cell 63 (4), 633–646 (2016).

38 Kim, S., Kim, D., Cho, S.W., Kim, J., & Kim, J.S., Highly efficient RNA-guided genome editing in human cells via delivery of purified Cas9 ribonucleoproteins. Genome research 24(6), 1012–1019 (2014).

39 Doench, J.G., Fusi, N., Sullender, M., Hegde, M., Vaimberg, E.W., Donovan, K.F., Smith, I., Tothova, Z., Wilen, C., Orchard, R., Virgin, H.W., Listgarten, J., & Root, D.E., Optimized sgRNA design to maximize activity and minimize off-target effects of CRISPR-Cas9. Nature biotechnology 34 (2), 184–191 (2016).

40 Kleinstiver, B.P., Pattanayak, V., Prew, M.S., Tsai, S.Q., Nguyen, N.T., Zheng, Z., & Joung, J.K., High-fidelity CRISPR-Cas9 nucleases with no detectable genome-wide off-target effects. Nature 529 (7587), 490–495 (2016).

41 Slaymaker, I.M., Gao, L., Zetsche, B., Scott, D.A., Yan, W.X., & Zhang, F., Rationally engineered Cas9 nucleases with improved specificity. Science 351 (6268), 84–88 (2016).

42 Fu, B.X., Hansen, L.L., Artiles, K.L., Nonet, M.L., & Fire, A.Z., Landscape of target:guide homology effects on Cas9-mediated cleavage. Nucleic acids research 42 (22), 13778–13787 (2014).

43 Horlbeck, M.A., Gilbert, L.A., Villalta, J.E., Adamson, B., Pak, R.A., Chen, Y., Fields, A.P., Park, C.Y., Corn, J.E., Kampmann, M., & Weissman, J.S., Compact and highly active next-generation libraries for CRISPR-mediated gene repression and activation. eLife 5 (2016).

44 Horlbeck, M.A., Witkowsky, L.B., Guglielmi, B., Replogle, J.M., Gilbert, L.A., Villalta, J.E., Torigoe, S.E., Tjian, R., & Weissman, J.S., Nucleosomes impede Cas9 access to DNA in vivo and in vitro. eLife 5 (2016).

45 Isaac, R.S., Jiang, F., Doudna, J.A., Lim, W.A., Narlikar, G.J., & Almeida, R., Nucleosome breathing and remodeling constrain CRISPR-Cas9 function. eLife 5 (2016).

46 Larson, M.H., Gilbert, L.A., Wang, X., Lim, W.A., Weissman, J.S., & Qi, L.S., CRISPR interference (CRISPRi) for sequence-specific control of gene expression. Nature protocols 8 (11), 2180–2196 (2013).

47 Gilbert, L.A., Horlbeck, M.A., Adamson, B., Villalta, J.E., Chen, Y., Whitehead, E.H., Guimaraes, C., Panning, B., Ploegh, H.L., Bassik, M.C., Qi, L.S., Kampmann, M., & Weissman, J.S., Genome-Scale CRISPR-Mediated Control of Gene Repression and Activation. Cell 159 (3), 647–661 (2014).

48 Perez-Pinera, P., Kocak, D.D., Vockley, C.M., Adler, A.F., Kabadi, A.M., Polstein, L.R., Thakore, P.I., Glass, K.A., Ousterout, D.G., Leong, K.W., Guilak, F., Crawford, G.E., Reddy, T.E., & Gersbach, C.A., RNA-guided gene activation by CRISPR-Cas9-based transcription factors. Nature methods 10 (10), 973–976 (2013).

49 Maeder, M.L., Linder, S.J., Cascio, V.M., Fu, Y., Ho, Q.H., & Joung, J.K., CRISPR RNA-guided activation of endogenous human genes. Nature methods 10 (10), 977–979 (2013).

50 Doench, J.G., Hartenian, E., Graham, D.B., Tothova, Z., Hegde, M., Smith, I., Sullender, M., Ebert, B.L., Xavier, R.J., & Root, D.E., Rational design of highly active sgRNAs for CRISPR-Cas9-mediated gene inactivation. Nature biotechnology 32 (12), 1262–1267 (2014).

51 Heigwer, F., Kerr, G., & Boutros, M., E-CRISP: fast CRISPR target site identification. Nature methods 11 (2), 122–123 (2014).

52 Haeussler, M., Schonig, K., Eckert, H., Eschstruth, A., Mianne, J., Renaud, J.B., Schneider-Maunoury, S., Shkumatava, A., Teboul, L., Kent, J., Joly, J.S., & Concordet, J.P., Evaluation of off-target and on-target scoring algorithms and integration into the guide RNA selection tool CRISPOR. Genome biology 17 (1), 148 (2016).

53 Haeussler, M. & Concordet, J.P., Genome Editing with CRISPR-Cas9: Can It Get Any Better? Journal of genetics and genomics = Yi chuan xue bao 43 (5), 239–250 (2016).

54 Hendel, A., Bak, R.O., Clark, J.T., Kennedy, A.B., Ryan, D.E., Roy, S., Steinfeld, I., Lunstad, B.D., Kaiser, R.J., Wilkens, A.B., Bacchetta, R., Tsalenko, A., Dellinger, D., Bruhn, L., & Porteus, M.H., Chemically modified guide RNAs enhance CRISPR-Cas genome editing in human primary cells. Nature biotechnology 33 (9), 985–989 (2015).

55 Shalem, O., Sanjana, N.E., Hartenian, E., Shi, X., Scott, D.A., Mikkelson, T., Heckl, D., Ebert, B.L., Root, D.E., Doench, J.G., & Zhang, F., Genome-scale CRISPR-Cas9 knockout screening in human cells. Science 343 (6166), 84–87 (2014).

56 Wang, T., Birsoy, K., Hughes, N.W., Krupczak, K.M., Post, Y., Wei, J.J., Lander, E.S., & Sabatini, D.M., Identification and characterization of essential genes in the human genome. Science 350 (6264), 1096–1101 (2015).

57 Blomen, V.A., Majek, P., Jae, L.T., Bigenzahn, J.W., Nieuwenhuis, J., Staring, J., Sacco, R., van Diemen, F.R., Olk, N., Stukalov, A., Marceau, C., Janssen, H., Carette, J.E., Bennett, K.L., Colinge, J., Superti-Furga, G., & Brummelkamp, T.R., Gene essentiality and synthetic lethality in haploid human cells. Science 350 (6264), 1092–1096 (2015).

58 Hart, T., Chandrashekhar, M., Aregger, M., Steinhart, Z., Brown, K.R., MacLeod, G., Mis, M., Zimmermann, M., Fradet-Turcotte, A., Sun, S., Mero, P., Dirks, P., Sidhu, S., Roth, F.P., Rissland, O.S., Durocher, D., Angers, S., & Moffat, J., High-Resolution CRISPR Screens Reveal Fitness Genes and Genotype-Specific Cancer Liabilities. Cell 163 (6), 1515–1526 (2015).

59 Munch, J., Rajan, D., Schindler, M., Specht, A., Rucker, E., Novembre, F.J., Nerrienet, E., Muller-Trutwin, M.C., Peeters, M., Hahn, B.H., & Kirchhoff, F., Nef-mediated enhancement of virion infectivity and stimulation of viral replication are fundamental properties of primate lentiviruses. Journal of virology 81 (24), 13852–13864 (2007).

60 Munk, C. & Landau, N.R., Production and use of HIV-1 luciferase reporter viruses. Current protocols in pharmacology Chapter 12, Unit12 15 (2003).

61 O’Doherty, U., Swiggard, W.J., & Malim, M.H., Human immunodeficiency virus type 1 spinoculation enhances infection through virus binding. Journal of virology 74 (21), 10074–10080 (2000).

62 Mandegar, M.A., Huebsch, N., Frolov, E.B., Shin, E., Truong, A., Olvera, M.P., Chan, A.H., Miyaoka, Y., Holmes, K., Spencer, C.I., Judge, L.M., Gordon, D.E., Eskildsen, T.V., Villalta, J.E., Horlbeck, M.A., Gilbert, L.A., Krogan, N.J., Sheikh, S.P., Weissman, J.S., Qi, L.S., So, P.L., & Conklin, B.R., CRISPR Interference Efficiently Induces Specific and Reversible Gene Silencing in Human iPSCs. Cell stem cell 18 (4), 541–553 (2016).

63 Brinkman, E.K., Chen, T., Amendola, M., & van Steensel, B., Easy quantitative assessment of genome editing by sequence trace decomposition. Nucleic acids research 42 (22), e168 (2014).

64 Koressaar, T. & Remm, M., Enhancements and modifications of primer design program Primer3. Bioinformatics 23 (10), 1289–1291 (2007).

65 Untergasser, A., Cutcutache, I., Koressaar, T., Ye, J., Faircloth, B.C., Remm, M., & Rozen, S.G., Primer3--new capabilities and interfaces. Nucleic acids research 40 (15), e115 (2012).

66 Wehrly, K. & Chesebro, B., p24 antigen capture assay for quantification of human immunodeficiency virus using readily available inexpensive reagents. Methods 12 (4), 288–293 (1997).

